# Development of the Olfactory and Vomeronasal Systems in the Fossorial Water Vole (*Arvicola scherman*). I. The Late Prenatal Stages

**DOI:** 10.1101/2025.08.30.673214

**Authors:** Sara Ruiz-Rubio, Irene Ortiz-Leal, Mateo V. Torres, Aitor Somoano, Taej¡kyun Shin, Pablo Sanchez-Quinteiro

## Abstract

Chemical communication is essential for mammalian survival from the earliest stages of life, yet most of what is known about the prenatal development of the olfactory and vomeronasal systems comes from laboratory rodents. These models, while invaluable, may not fully represent the developmental trajectories of wild species living under natural ecological pressures. Here we investigated the fetal development of the nasal chemosensory systems in the fossorial water vole (*Arvicola scherman*), a free-living arvicoline rodent with a highly subterranean lifestyle. We analyzed fetuses at embryonic days E17 and E21 (term) using classical histology, immunohistochemistry (markers: Gαi2, Gαo, Gγ8, CB, CR, PGP 9.5, GAP-43, β-tubulin, MAP2), and lectin histochemistry (UEA, LEA, SBA, STA, DBA). This combined approach enabled us to assess structural maturation, neuronal differentiation, and the temporal dynamics of glycoconjugate expression in the vomeronasal organ (VNO), olfactory epithelium (OE), and the main (MOB) and accessory olfactory bulbs (AOB).

By E21, the MOB displayed a six-layered adult-like organization with well-defined glomeruli and interneuronal populations, whereas the AOB showed delayed morphological maturation but already exhibited selective molecular signatures in its nerve and superficial layers. Prenatally, the VNO underwent conspicuous structural differentiation, including stratification of the sensory epithelium, robust axonal fasciculation, and early development of vomeronasal glands. Immunohistochemical analysis revealed early expression of G-protein subunits and calcium-binding proteins, indicating premature pathway specification and interneuronal circuit formation. Lectin labeling provided additional insights: SBA emerged as a highly selective marker of the vomeronasal pathway; UEA highlighted early compartmentalization of vomeronasal projections; LEA showed a conserved, pan-chemosensory binding pattern across systems; and DBA, despite its lower specificity, revealed late-onset reactivity in postmitotic neurons. Together, these findings demonstrate that *A. scherman* exhibits a remarkably accelerated prenatal maturation of its chemosensory systems compared with laboratory rodents. This early functional readiness likely reflects adaptive pressures of a fossorial lifestyle, emphasizing the importance of incorporating wild species into developmental neurobiology to refine our understanding of mammalian chemosensory evolution.

## Introduction

In mammals, chemical communication is critical for survival from the earliest stages of life (Surov and Maltsev 2016). Far beyond its role in adult social and reproductive behaviors, chemical signaling guides crucial perinatal interactions, including maternal recognition, initiation of suckling, sibling discrimination, and early territorial imprinting (Dulac and Torello 2003; Coria-Avila et al. 2022). From a developmental perspective, the timing of sensory-neuron differentiation and the early assembly of bulb circuits constrain chemosensory competence at birth, shaping neonatal fitness trajectories. Among the olfactory subsystems, the vomeronasal system (VNS) plays a particularly central role in mediating innate responses to pheromonal and kairomonal cues, often triggering hardwired behaviors essential for neonatal viability and long-term species propagation (Liberles 2014; Torres et al. 2023b; Sanmartín-Vázquez et al. 2024).

The VNS comprises the vomeronasal organ (VNO), the accessory olfactory bulb (AOB), and a well-defined array of central projections to the amygdala and hypothalamus (Scalia and Winans 1975; Salazar et al. 2016; Mohrhardt et al. 2018). The sensory epithelium of the VNO houses receptor neurons that express vomeronasal receptors (V1R, V2R, and FPRs), which recognize social and environmental chemical cues (Rivière et al. 2009; Torres et al. 2021; Ortiz-Leal et al. 2024). These neurons project to the AOB, where signals undergo initial processing before being relayed to higher-order brain centers. In most mammals with a well-developed vomeronasal system—such as rodents (Enomoto et al. 2011), marsupials (Torres et al. 2022), and lagomorphs (Villamayor et al. 2020)—vomeronasal projections segregate into two distinct zones within the AOB: an anterior region receiving inputs from V1R-expressing neurons and a posterior region connected to V2R-expressing neurons (Rodriguez et al. 1999; Halpern and Martínez-Marcos 2003).

However, this segregated model is not ubiquitous among mammals. In major groups within Laurasiatheria (Carnivora (Salazar et al. 1998), Cetartiodactyla (Kondoh et al. 2017a), Perissodactyla (Chun et al. 2024), and Soricidae (Oikawa et al. 1993)), there has been a transition toward a uniform model characterized by the absence of V2R projections to the AOB, likely due to the pseudogenization of the corresponding receptors (Grus and Zhang 2009; Prieto-Godino et al. 2016; Policarpo et al. 2024). This loss remains controversial, as some observations in artiodactyls (e.g., pigs and cows) (Kondoh et al. 2022) and wild canids (Ortiz-Leal et al. 2022b; Ortiz-Leal et al. 2024) suggest the presence of functional V2R genes and hypothetical projections to the transitional zone between the MOB and the AOB, known as the olfactory limbus (Larriva-Sahd 2012; Ortiz-Leal et al. 2023; Ortiz-Leal et al. 2024). Regardless of model, the vomeronasal pathway governs an array of behaviors—from mating and aggression to predator avoidance—making it an indispensable part of the mammalian chemosensory repertoire (Brennan and Zufall 2006). These macroevolutionary departures underscore that organizational principles inferred from a few model species may not universally apply, especially when ecological pressures differ.

Despite this significance, most of what we know about prenatal and early postnatal development of the VNS (and olfactory systems more broadly) derives from laboratory mouse and rat (Katreddi et al. 2022; Tufo et al. 2022; Kim et al. 2023), whose domestication, captive environments, and reduced genetic variability can limit ecological generalization (Würbel 2000; Richter et al. 2009; Modlinska and Pisula 2020; Savriama et al. 2022). To better assess the generalizability of laboratory-derived patterns, it is important to incorporate evidence from alternative models in wild, non-domesticated rodents. Examining free-living species that experience natural sensory ecologies, where selective pressures on chemosensation—predation risk, subterranean life, diet, and social organization— differ markedly from laboratory conditions, will help distinguish core developmental features from shifts driven by artificial selection. In this context, the fossorial water vole *Arvicola scherman*—also considered as a fossorial form of *A. amphibius* (Balmori-de La Puente et al. 2022)—stands out as an ecologically and economically important model. This cricetid rodent occupies extensive burrow systems in meadows, grasslands, and fruit orchards from lowlands to elevations of ∼2000 m a.s.l., across the northern Iberian Peninsula, the Alps, central European mountains, and the Carpathians (Musser and Carleton 2005; Somoano 2024). It shows a strongly fossorial lifestyle, feeding mainly on roots, bulbs, and tubers of dicotyledons and some Poaceae (Airoldi 1976). Due to their high reproductive output (Somoano et al. 2016; Somoano et al. 2017) and capacity to reach population densities exceeding 1,000 individuals/ha during population outbreaks, they are considered major agricultural pests (Giraudoux 1997). Furthermore, *A. scherman* is a known reservoir of zoonotic pathogens including *Leptospira* spp., *Borrelia burgdorferi*, *Echinococcus multilocularis* and *Toxoplasma gondii*, posing additional risks to public health (Viel et al. 1999; Fuehrer et al. 2010; Espí et al. 2017).

Given its fossorial nature, *A. scherman* has evolved a strong reliance on olfactory and vomeronasal cues to compensate for reduced visual and auditory stimuli in subterranean environments (Dennis et al. 2020). This ecological trait renders the study of its chemosensory systems particularly insightful, especially regarding the functional maturation of the VNS in prenatal stages—a subject hitherto unaddressed. Our previous studies have provided a comprehensive characterization of the adult VNO and AOB in this species using histological, immunohistochemical, and lectin-histochemical methods (Ruiz-Rubio et al. 2024a; Ruiz-Rubio et al. 2024b).

In this work, we have investigated the fetal development of the nasal chemosensory systems of *A. scherman*, with emphasis on the VNO and AOB, to assess how far laboratory-derived developmental hallmarks extend to a fossorial, free-living arvicoline. We have focused on two fetal ages: E17, which in the laboratory mouse aligns with the onset of axonal invasion of the presumptive glomerular layer and the initial emergence of protoglomerular units (≈E17–E18), and E21, which corresponds to the at-term stage in arvicoline voles (gestation ≈ 21 days) (Quéré 2009). This sampling design allowed us to contrast, in *A. scherman*, the initiation of glomerular assembly with its at-term state. Specifically, we examined (i) olfactory and vomeronasal glomerulogenesis, including the coordinated emergence of periglomerular and granule-cell interneurons that populate the glomerular and granule layers, the timing of neuronal maturation markers—OMP, tubulin, PGP9.5— and (iii) the onset of vomeronasal specification associated with G-protein pathways (Gαi2 and Gαo). We employed classical histology, together with immunohistochemistry and lectin histochemistry, to characterize this structural maturation and molecular specialization. This comparative approach clarifies which elements of the laboratory paradigm are conserved, shifted, or divergent under natural sensory ecologies. Additionally, to our knowledge, this is the first study to document the ontogeny of the vomeronasal system in a wild cricetidae species.

## Methods

### Specimen collection

This study was carried out on fetal specimens of the fossorial water vole (*Arvicola scherman*). As part of the integrated control program (Article 15.127 of Spanish Law 43/2002 on Plant Health) in the province of Lugo (Galicia), trapping was conducted in meadowlands surrounding the village of Triacastela. Snap traps (Supercat®, Swissinno, Switzerland) were placed in active burrow systems and left operational for 24 h. These devices typically induce immediate death by head trauma; when this did not occur, cervical dislocation was promptly performed in accordance with Directive 2010/63/EU on the protection of animals used for scientific purposes. In addition, litters from two females were incidentally obtained during live-trapping campaigns conducted to monitor population dynamics.

The first litter consisted of embryos with a crown–rump length (CRL) of 4,2 cm, while the second included smaller fetuses of 3,1 cm CRL, corresponding to earlier developmental stages (Fig. 1). The presence of the fetuses was discovered post-mortem during routine necropsy procedures. In a previous study, as part of the integrated control program for the fossorial water vole in the province of Lugo, a necropsy was conducted on 194 female specimens previously captured using snap traps. Of these, 49 were pregnant. Embryos extracted from their uteri were measured for crown-rump length (CRL) with a digital caliper, ensuring a precision of 0.01 mm. Recorded lengths ranged from 3.80 mm to 42.00 mm. To estimate embryonic age in days post-coitum (DPC), available data on embryonic development from *Mus musculus*, which, to our knowledge, represents the closest phylogenetic relative with a comparable gestation period, was used as a reference to determine DPC at the minimum observed CRL (Graham et al., 2015). Given that the gestation period of *A. scherman* spans 21 to 22 days (Quéré, 2009) the maximum embryonic length was assigned to near-term fetuses (21 days), while the minimum length corresponded to embryos first visible to the naked eye (10 days). To refine these estimations, a growth regression model was applied using IBM SPSS Statistics 20.0, with CRL as the independent variable and known DPC values as the dependent variable. The resulting regression equation provided predicted DPC values for embryos with unknown age, enhancing the accuracy of developmental assessments. In conclusion, the litters identified with a CRL of 31 mm and 42 mm corresponded to developmental stages of 16.96 days (E17) and 21.0 days (E21), respectively.

**Figure 1.**
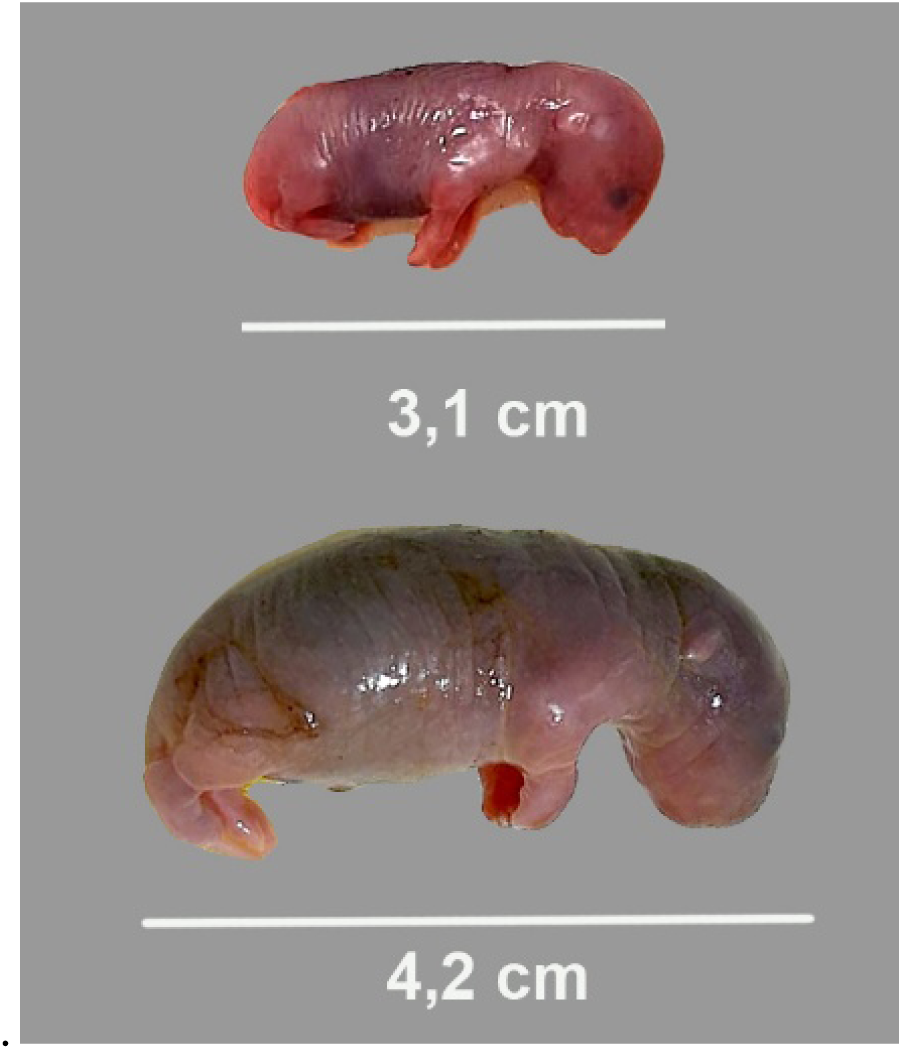
Representative examples of two fetuses from the two litters used in the present study. The first fetus (3.1 cm CR length) corresponds to embryonic day 17 (E17), while the second one (4.2 cm CR length) corresponds to embryonic day 21 (E21). The crown-rump (CR) length is indicated below each specimen.

All fetuses were immediately harvested and immersed in freshly prepared Bouin’s solution to ensure optimal fixation. Transportation of the samples from the capture site to the laboratory never exceeded four hours, minimizing the risk of tissue degradation

### Dissection and anatomical processing

Due to the delicate condition of the fetal specimens and the incomplete ossification of cranial structures at these developmental stages, tissue handling was performed under a stereomicroscope using fine microsurgical instruments. In most cases, the mandibles were first removed. After 24 hours of fixation in Bouin’s fluid, the heads were removed and, without further manipulation, post-fixed in 70% ethanol until paraffin embedding. Due to the absence of significant mineralization, decalcification was not required, which facilitated optimal antigen preservation.

### Processing samples for microscopic study

Tissues were processed through a graded ethanol dehydration series (70%, 90%, 96%, and absolute ethanol), followed by clearing in a 1:1 ethanol–xylene solution and two changes of pure xylene. Paraffin infiltration was performed at 60°C for 3 hours. Once embedded, paraffin blocks were trimmed and sectioned using a rotary microtome. Sections were cut at 6–8 µm thickness in the transverse plane. Serial sections were mounted on glass slides and dried overnight at 37°C.

### General histological staining

Nissl staining with 1% cresyl violet allowed structural analysis of the nasal cavity and neuronal profiling within the developing VNOs and olfactory bulbs.

### Immunohistochemical staining

Immunohistochemical staining was carried out to characterize cell types, neuronal maturation, and the expression of signaling proteins involved in vomeronasal and olfactory transduction. Sections were first deparaffinized and rehydrated, followed by quenching of endogenous peroxidase activity with 3% hydrogen peroxide in distilled water. Non-specific binding was blocked using 2% bovine seroalbumin (BSA). After overnight incubation with the primary antibodies at 4°C, samples were washed three times in PB and then incubated for 30 min with the CRF Anti-Polyvalent HRP Polymer (ScyTek, Cache Valley, USA). Signal visualization was achieved using 3,3-diaminobenzidine (DAB) in 0.2 M Tris-HCl buffer (pH 7.61) containing 0.003% H₂O₂. Slides were counterstained with Mayer’s hematoxylin, dehydrated, and mounted. Negative controls included omission of the primary antibody.

The immunohistochemical analysis included the following primary antibodies that provided valuable insights into the morphofunctional characteristics of the VNS. More detailed information is provided in Table 1.

**Table 1.** Detailed information on the antibodies used in this study: Species of elaboration, dilution, catalogue number, manufacturer and target immunogens.

| Antibody | 1st Ab species | Dilution | Supplier / Catalog number | Target immunogen |
| --- | --- | --- | --- | --- |
| Anti-Gao | Rabbit | 1:200 | MBL-551 | Bovine GTP binding protein Gao subunit |
| Anti-Gai2 | Rabbit | 1:100 | Santa Cruz Biotech. SC-7276 | Peptide mapping within a highly divergent domain of Gai2 of rat origin |
| Anti-Gg8 | Rabbit | 1:200 | Cloud-Clone PAQ769Mu01 | Recombinant G Protein Gamma 8 (GNg8) |
| Anti-CB | Rabbit | 1:200 | Proteintech 14479-1-AP | Recombinant CB expressed in *E. coli*. |
| Anti-CR | Rabbit | 1:200 | Proteintech 12278-1-AP | Recombinant human CR containing an N-terminal 6×His (hexahistidine) tag |
| Anti-MAP-2 | Mouse | 1:200 | Sigma M4403 | Rat brain microtubule-associated proteins |
| Anti-GAP-43 | Rabbit | 1:200 | Proteintech 16971-1-AP | GAP43 fusion protein Ag9294 |
| Anti-TUB | Rabbit | 1:300 | Proteintech 10068-1-AP | β-Tubulin fusion protein Ag0117 |
| Anti-PGP | Rabbit | 1:200 | Proteintech 14730-1-AP | UCHL1/PGP9.5 fusion protein Ag6490 |
| Abbreviations: CB, calbindin; CR, calretinin; GAP-43, growth-associated protein-43; MAP-2, microtubule-associated protein-2; PGP 9.5, protein gene product 9.5; TUB, β-tubulin class III. |  |  |  |  |

#### G-protein subunits

Anti-Gαi2: Recognizes the Gαi2 subunit, a key component of the intracellular transduction cascade associated with V1R-type vomeronasal receptors (Torres et al., 2024).

Anti-Gαo: Binds to the Gαo subunit, which participates in the signal transduction mechanism downstream of V2R-type receptors (Villamayor et al., 2021).

Anti-Gg8: G-protein expressed in the developing axons of olfactory and vomeronasal neurons (Ryba and Tirindelli, 1995; Tirindelli and Ryba, 1996).

#### Neuronal differentiation and plasticity

Anti-Growth-Associated Protein 43 (GAP-43): Indicator of neuronal growth and axonal pathfinding (Benowitz and Routtenberg, 1997).

#### Cytoskeletal proteins in projection neurons

Anti-Microtubule-associated-protein-2 (MAP-2): Microtubular component in neuronal dendrites (Dehmelt and Halpain, 2005).

Anti-β-tubulin class III (TUB) : Neuron-specific cytoskeletal protein used as a pan-neuronal marker during development. It highlights neuronal architecture and early differentiation (Menezes and Luskin, 1994).

#### Calcium-binding proteins

Anti-Calbindin-D28k (CB): Involved in intracellular calcium buffering. CB is a useful marker for delineating specific neuronal populations in sensory pathways (Andressen et al., 1993).

Anti-Calretinin (CR): Expressed in distinct neuronal subsets, often complementing calbindin distribution. It serves as a reliable marker for interneurons and sensory relay cells (Jacobowitz and Winsky, 1991).

Anti-PGP 9.5: Also known as UCHL1, this ubiquitin C-terminal hydrolase is widely expressed in neurons and neuroendocrine cells. It is routinely used as a pan-neuronal marker to highlight general neuronal distribution across central and peripheral tissues (Thompson et al., 1983).

### Lectin histochemical labelling

Lectin labelling, while similar to immunohistochemical techniques, relies on specific proteins known as lectins. These proteins have domains that recognize and bind non-covalently to terminal sugars in tissues, forming glycoconjugates. Unlike antibodies, lectins do not have an immune origin (Lis and Sharon, 1998). Lectins have been extensively used in the study of the olfactory systems (Shin et al., 2017). In this study, we employed five lectins. More details in Table 2.

**Table 2.** Lectins used, concentrations and binding specificities.

| Lectin | Abbrev. | Dilut. mg/ml | Supplier Catalog number | Preferred sugar specificity | Specificity groups |
| --- | --- | --- | --- | --- | --- |
| *Ulex europaeus* (Gorse) agglutinin | UEA | 2.0 | Vector B-1065-2 | α-Fuc | Fucose |
| *Lycopersicon esculentum* (Tomato) lectin | LEA | 1.0 | Vector B-1175-1 | β-1,4 GlcNAc oligomers | GlcNAc |
| *Glycine max* (Soybean) agglutinin | SBA | 2.0 | Vector 1015-5 | GalNAc | Gal/GalNAc |
| *Solanum tuberosum* (Potato) lectin | STA | 2.0 | Vector B-1165-2 | GlcNAc Oligomers, LacNAc | GlcNAc |
| *Dolichos biflorus* (Horse gram) lectin | DBA | 2.0 | Vector B-1035-5 | $\alpha$ GalNAc | GalNAc |
| Abbreviations: Fuc, fucose; Gal, galactose; GalNAc, N-acetyl-galactosamine; GlcNAc, N-acetyl-glucosamine; LacNAc, N-acetyl-lactosamine. |  |  |  |  |  |

#### Ulex europaeus (UEA)

UEA specifically binds to terminal L-fucose present in glycoproteins and glycolipids. It was chosen for this study because of its role as a specific marker for the AOB in several species, including mice (Kondoh et al., 2017a), fox (Ortiz-Leal et al., 2022), and wolves (Ortiz-Leal et al., 2024).

#### Lycopersicum esculentum (LEA)

LEA exhibits high affinity for N-acetylglucosamine and has been widely used as a comprehensive marker for both the main and accessory olfactory bulb in various species. These include rodents such as rats (Franceschini et al., 1994), mice (Keller et al., 2022), capybaras (Torres et al., 2020), and hamster (Taniguchi et al., 1993b).

#### Glycine max (SBA)

SBA, isolated from soybean, binds specifically to N-acetylgalactosamine residues. SBA binds to vomeronasal sensory neurons and the accessory olfactory bulb in goats (Park et al., 2014).

#### Solanum tuberosum (STA)

STA, which is isolated from potato, binds oligomers of N-acetylglucosamine and some oligosaccharides containing N-acetylglucosamine and N-acetylmuramic acid (Chun et al., 2024). This lectin has previously been used in the characterization of the VNS in horses, bears, and dama gazelles (Lee et al., 2016; Tomiyasu et al., 2018; Torres et al., 2023a).

#### Dolichos biflorus (DBA)

DBA is isolated from horse gram and has a carbohydrate specificity toward N-acetylgalactosamine (Hofmann and Meyer, 1991). DBA is an excellent marker for AOB neurons in species such as goat and sheep (Mogi et al., 2007; Salazar et al., 2000).

The lectin histochemical labelling process began with slides that had been deparaffinized and rehydrated. To suppress endogenous peroxidase activity, they were treated with a 3% H_2_O_2_ solution. A 2% bovine serum albumin (BSA) solution was applied to block non-specific bindings. The lectins were incubated overnight at 4°C. Following this, the slides were exposed to a 90-minute incubation in an avidin-biotin complex (ABC) solution with peroxidase (Vectorlabs, Burlingame, CA). This complex interacts with the lectin during the incubation, enhancing the subsequent peroxidase reaction. The reaction was visualized using a solution of 0.003% H_2_O_2_ and 0.05% 3,3-diaminobenzidine (DAB) in 0.2 M Tris-HCl buffer, which produced a brown precipitate. Controls included tests without the lectin, as well as with lectins pre-absorbed with an excess of their corresponding sugars.

### Image acquisition

Digital images were acquired using an Olympus SC180 camera attached to an Olympus BX50 microscope. To create composite images from multiple photographs, PTGuiPro software (Rotterdam, Netherlands) was employed due to the large size of the studied structures. Adobe Photoshop CS4 (San Jose, CA) was utilized to modify brightness, contrast, and white balance levels; however, no enhancements, additions, or alterations to the image characteristics were performed.

Following photomicrography of the immune- and lectin-labeled sections, coverslips were carefully removed by immersion in xylene for approximately 48 hours. The sections were then rehydrated through a graded series of alcohols and distilled water, counterstained with hematoxylin, and subsequently dehydrated, cleared, and coverslipped again. This procedure allowed for enhanced visualization of the specific localization of the immunolabeling.

## Results

### Histological study

The application of Nissl staining enabled a precise histological identification of the main structural components of the vomeronasal organ, the olfactory mucosa, and the associated rostral cranial structures in the E17 *Arvicola scherman* fetus (Fig. 2). At a transverse level through the central portion of the head, the paired vomeronasal organs are clearly visible at the base of the nasal septum, embedded within the surrounding cartilaginous structures and showing early differentiation of the VNO (Fig. 2A).

**Figure 2.**
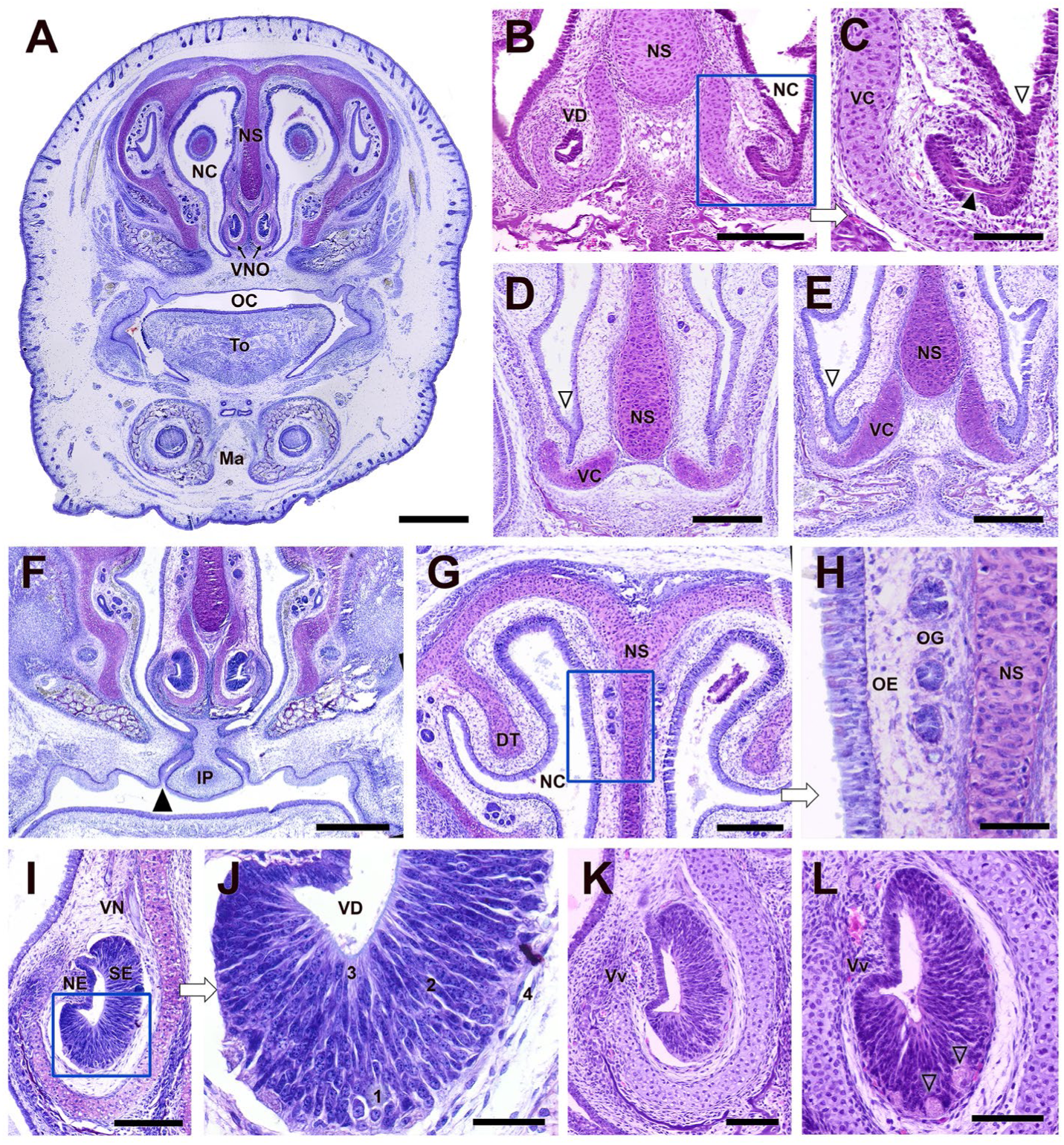
Histological study of the vomeronasal organ (VNO) and olfactory epithelium (OE) in the E17 embryo of the fossorial water vole. **(A)** Transverse section of the whole head at the mid-VNO level showing the paired VNOs located at the base of the nasal septum (NS), within the nasal cavity (NC). The VNO sensory epithelium (SE) is neatly distinguishable. **(B– E)**. Rostral portion of the nasal septum in two specimens. The most anterior part of the vomeronasal duct (VD; black arrowhead) and the ventral recess of the NS (open arrowhead) can be observed. The VNO cartilage (VC) initially shows a V-shape (D) and evolves in a more caudal level toward the characteristic J-shape (E). **(F)** Section at the level of the incisive papilla (IP; black arrowhead), showing its topographical relationship with surrounding structures and its potential role in allowing fluid or molecule passage from the oral to the nasal cavity (NC). **(G–H)** Olfactory epithelium (OE) possesses a reduced cellularity. Prominent olfactory glands (OG) are visible in the lamina propria. **(I–L)** Detailed view of the vomeronasal duct epithelium (VD), showing a stratified structure with large, rounded nuclei (1), intermediate fusiform cells (2–3), and flattened apical cells (4). Thick vomeronasal nerve (VN) bundles are present in a dorsomedial location. No glandular structures are present. In more caudal sections (K, L), large nerve bundles in the ventral part of the SE and a small vomeronasal vein (Vv) in the lateral part of the VNO. Nissl staining. Abbreviations: DT, dorsal turbinate; Ma, mandible; To, tongue. Scale bars: (A,F) = 500 µm, (B,D,E,G,I) = 250 µm, (C,H,L) = 100 µm, (J,K) = 50 µm.

In the most rostral part of the nasal septum, it is possible to identify the most anterior segment of the vomeronasal duct, which appears as a narrow epithelial tube with a small lumen. Just beneath it, a well-defined recess emerges in the ventral part of the nasal septum (Fig. 2B,C); this structure is expected to establish the future communication between the nasal cavity and the vomeronasal duct. The rostral part of the vomeronasal cartilage undergoes a morphological transition from a V-shaped profile in the more anterior sections (Fig. 2D) to a characteristic J-shape at more caudal levels (Fig. 2E). At this stage of development, the incisive papilla is well differentiated (Fig. 2F). Its topographical arrangement highlights the anatomical connection between the oral and nasal cavities. This configuration suggests a functional pathway through which fluids or chemical signals may pass from the oral cavity to the vomeronasal system, even during fetal development.

The olfactory epithelium at this developmental stage is characterized by low cellular density and signs of early differentiation. The neuroepithelium shows a pseudostratified organization with cells at various stages of maturation. In the lamina propria, olfactory tubular glands are already prominent, suggesting the beginning of glandular function linked to the olfactory process (Fig. 2G,H).

The vomeronasal epithelium presents a multilayered architecture with distinct cellular compartments (Fig. 2I,J). The basal zone contains cells with large, rounded nuclei with dark nucleoli and pale cytoplasm. Intermediate cells progressively elongate and acquire a more fusiform morphology, while the apical region is occupied by flattened cells with dense chromatin. Basally positioned cells are also distinguishable. In the dorsomedial portion of the epithelium, vomeronasal nerve bundles are clearly observed. Unlike the olfactory mucosa, no glandular elements are present within the underlying lamina propria of the vomeronasal epithelium (Figures 2I,J). In more caudal sections, thick bundles of nerve fibers appear at the base of the sensory epithelium, along with a small-caliber lateral vomeronasal vein (Fig. 2L).

The Nissl-stained histological study of near-term E20 embryos reveals a more advanced stage of differentiation across all anatomical components of the vomeronasal and olfactory systems (Fig. 3). One of the most striking features is the presence at his stage of a well-defined connection between the nasal cavity and the vomeronasal duct, clearly identifiable at this stage (Fig. 3A). At this level, the vomeronasal cartilage forms a broad, symmetrical capsule that flanks the base of the nasal septum. The VNO displays an increased degree of structural maturation. The sensory epithelium maintains a comparable cellular density to earlier stages, but exhibits a more ordered laminar architecture, with clearly stratified cell layers and an apical zone devoid of nerve endings (Fig. 3B,C,F-H). In contrast, the nonsensory epithelium has expanded significantly and now comprises a larger proportion of the ductal lining (Fig. 3G,H).

**Figure 3.**
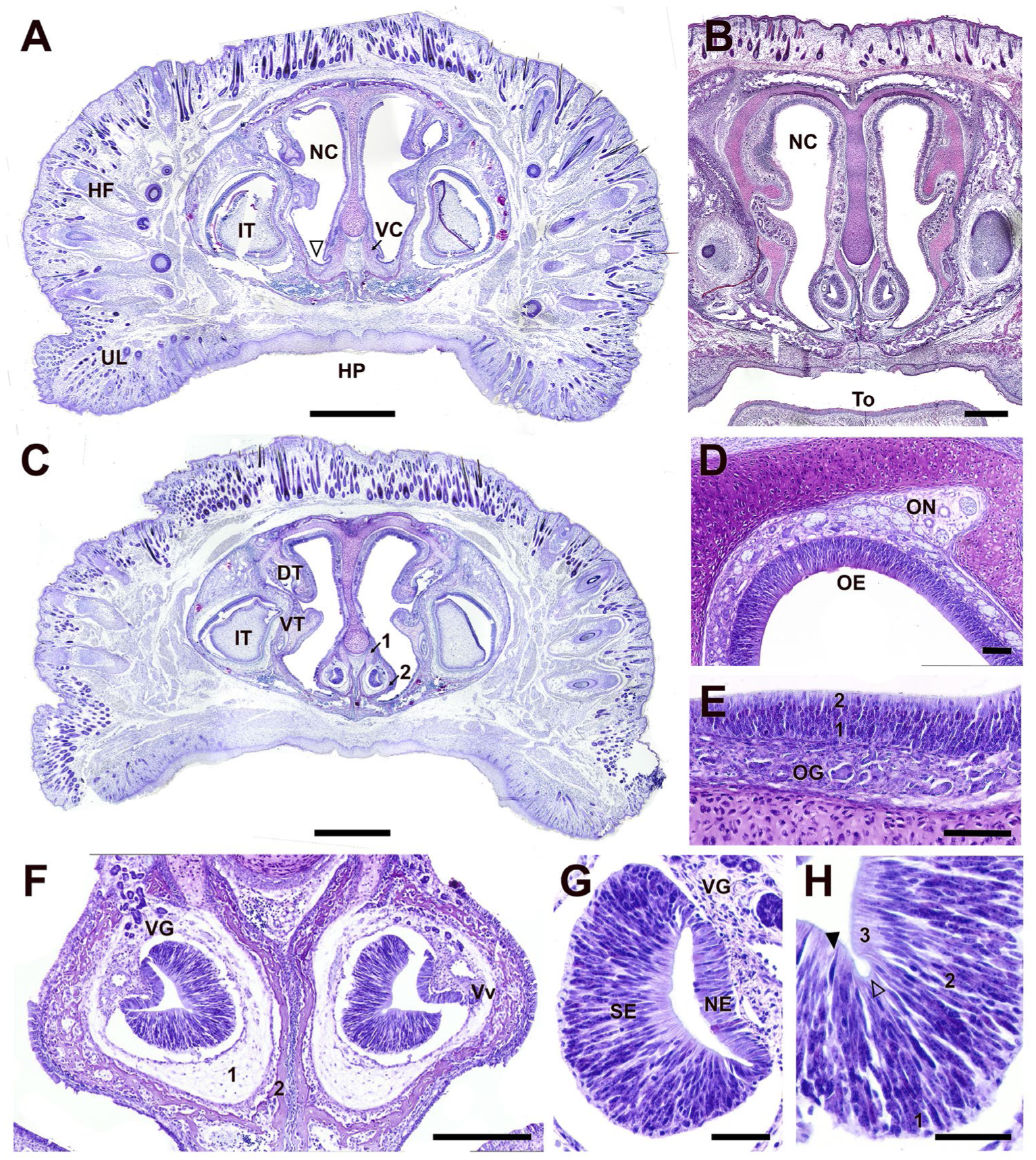
Nissl histological staining study of the VNO and OE in the E21 embryo of the fossorial water vole. **(A)** Transverse section at the level of communication between the nasal cavity and the opening of the vomeronasal organ (open arrowhead). At this level, the vomeronasal cartilage (VC) appears as a broad and well-defined capsule, bilaterally flanking the base of the nasal cavity (NC). **(B)** Section through a central region of the VNO shows the kidney-shaped profile of the VNO and the remarkable development of the sensory epithelium. **(C)** At the level of the incisor teeth (IT), the cartilage capsule (1), dominant in earlier stages, is now being progressively replaced by a thin bony capsule (2). **(D–E)** The OE shows a high cellular density, with a layered structure composed of a compact zone of densely packed nuclei (1) and a more superficial zone with loosely arranged individual cells (2). **(F)** Detailed view of the vomeronasal area showing partial dissolution of the vomeronasal cartilage (1) and the formation of the surrounding bony capsule (2). A dorsolateral opening in the capsule contains numerous tubular vomeronasal glands (VG). The vomeronasal vein (Vv) is evident. **(G–H)** High-magnification images of the vomeronasal epithelium. The sensory epithelium (SE) and non-sensory epithelium (NE) are distinguishable. Apical layers contain densely stained flattened nuclei (3), whereas intermediate cells (2) and basal cells are clearly layered. Tubular glands are also evident. Abbreviations: DT, dorsal turbinate; HF, hair follicle; HP, hard palate; OG, olfactory glands; ON, olfactory nerves; To, tongue; UL, upper lip; VT, ventral turbinate. Scale bars: (A,C) = 1 mm, (B) = 500 µm, (F) = 250 µm, (D,E) = 100 µm, (G,H) = 50 µm.

A remarkable feature is the progressive replacement of the vomeronasal cartilage by a thin layer of bone, forming a nascent bony capsule that surrounds the vomeronasal duct. This structure, originating from the vomer bone primordy, begins to encapsulate the organ completely and is especially evident near the incisor region (Fig. 3C). In this same area, the degenerated cartilage and newly forming bone can be distinguished, highlighting the ongoing remodeling of the vomeronasal sheath (Fig. 3F). A dorsolateral opening is visible within this capsule, containing well-formed tubular vomeronasal glands, which were not prominent at earlier stages (Fig. 3F,G).

The olfactory epithelium has undergone substantial reorganization. It displays a high and uniform cellular density, with a compact basal zone and more loosely arranged superficial layers, indicating progressive stratification (Fig. 3D,E). The lamina propria is densely populated with olfactory nerve bundles, and olfactory glands are prominently distributed throughout, reflecting functional maturation (Fig. 3E).

The histological study of the main and accessory olfactory bulbs (MOB and AOB) in fetal *Arvicola scherman*, using Nissl staining, reveals a progressive degree of differentiation between embryonic days 17 (E17) (Fig. 4A-F) and 21 (E21) (Fig. 4G-L). At E17, the MOB already displays a rudimentary organization, with identifiable zones including a superficial olfactory nerve layer, an emerging external plexiform layer, a compact band of principal cells (future mitral cells), and a forming granule cell layer in the core (Fig. 4C,F). However, the laminar architecture remains diffuse and olfactory glomeruli are not yet evident. The AOB, positioned dorsocaudally to the MOB, is present but exhibits a much less defined organization. Principal cells are loosely distributed and show a topographical alignment with the PC layer of the MOB (Fig. 4D,E). Despite its early appearance, the AOB at this stage lacks distinct layering and glomerular structures.

**Figure 4.**
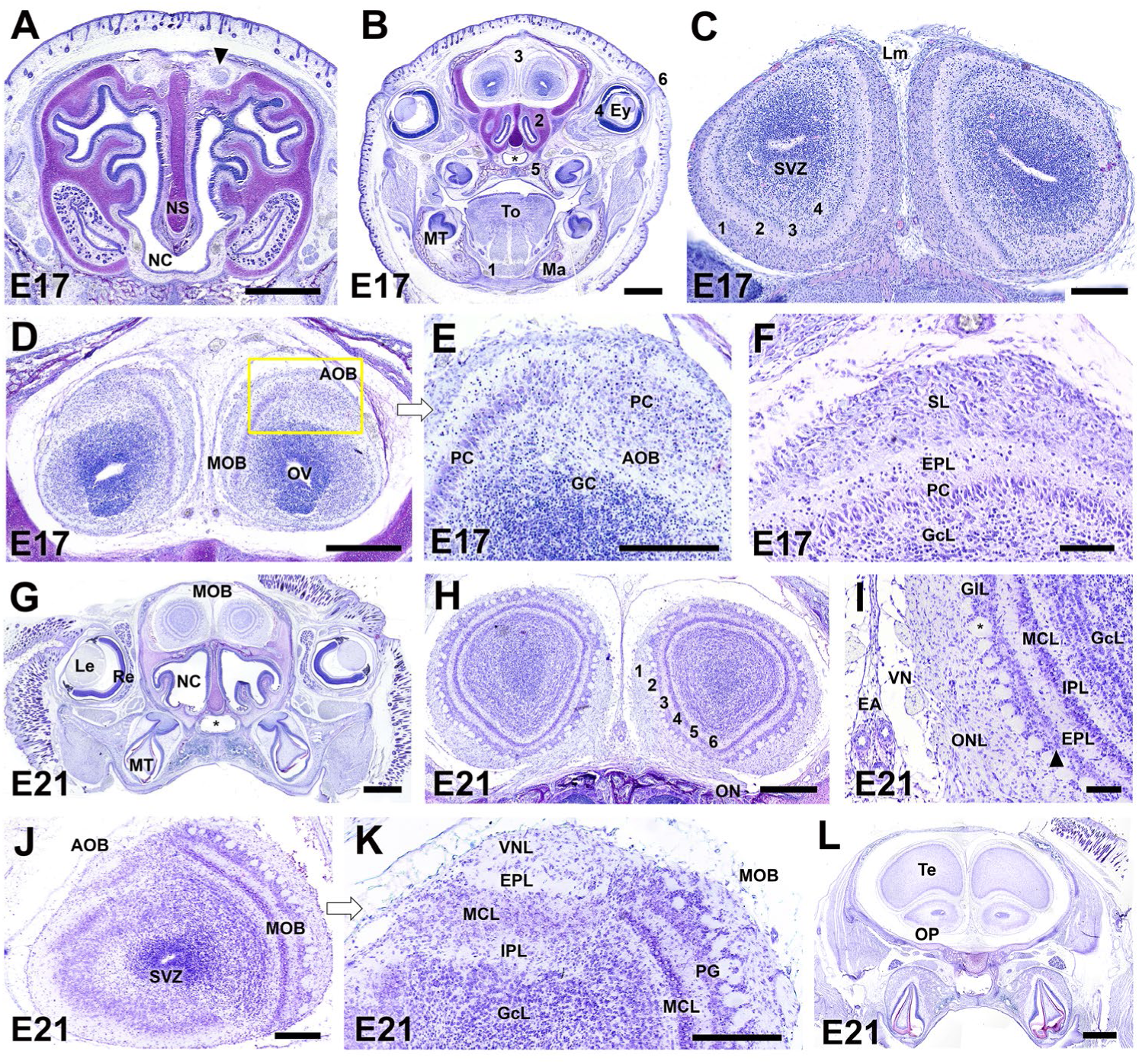
Histological study using Nissl staining of the main (MOB) and accessory (AOB) olfactory bulbs of the fossorial water vole at embryonic days 17 (A-F) and 21 (G-L). **(A)** Transverse section of the nasal cavity at the rostral level of the ethmoidal fossa. The anterior-most portion of the MOB is visible dorsally to the NS. **(B)** Section at the level of the eyes showing the caudal portion of the MOB and the AOB in a dorsal position. 1, Mylohyoid muscle; 2, Caudal recess of the nasal cavity; 3, OB; 4, Retina; 5, Palatine bone; 6, Palpebral fissure. **(C)** Central portion of the MOB displaying incipient lamination: from superficial to deep, four layers can be distinguished—(1) the olfactory nerve layer (ONL), (2) an external plexiform layer (EPL), (3) a dense band of principal cells (PC, primordial mitral cells), and (4) an internal plexiform layer overlying a developing granule cell layer (GcL). **(D,E)** Study of the AOB with a higher magnification of the box marked in D. The AOB comprises a diffuse cluster of principal cells, loosely organized but topographically aligned with the better-differentiated PC layer of the MOB. **(F)** The MOB at this stage reveals more clearly defined lamination than the AOB, with an external superficial layer (SL) lacking identifiable olfactory glomeruli, an external plexiform layer (EPL), a developing mitral cell layer (PC), and a deep granule cell layer (GcL). **(G-I)** At E21, the MOB presents a high degree of maturation. (G) Transverse section showing both MOBs and the nasal cavity, with well-developed surrounding structures. (H) The MOB reveals an adult-like laminated structure with six identifiable layers: (1) olfactory nerve layer (ONL), (2) glomerular layer (GL), (3) external plexiform layer (EPL), (4) mitral cell layer (MCL), (5) internal plexiform layer (IPL), and (6) a dense granule cell layer (GcL). (I) The glomeruli (asterisk) appear as discrete units aligned in a single layer, each surrounded by numerous periglomerular cells. Mitral cells are well-defined and flanked by the EPL and IPL. Medial to each MOB, the vomeronasal nerve (VN) bundles are clearly visible, highlighting the pathway of vomeronasal input. **(J–K)** The AOB remains less laminated than the MOB, though a vomeronasal nerve layer (VNL) is present. Glomeruli are absent. A dense band of principal cells with mitral-like morphology is seen (MCL), surrounded by less defined EPL and IPL layers. **(L)** A more caudal section shows the topographical relationship between the telencephalon (Te), olfactory peduncle (OP), and NC. Abbreviations: EA: Ethmoidal artery, Ey: Eyeball, Le: lens, Lm: leptomeninges, Ma: mandible, MT: Molar teeth, ON: Olfactory nerve, OV: Olfactory ventricle, Re: Retina, SVZ: Subventricular zone, Te: Telencephalon, To: Tongue. Scale bars: (A,B,G) = 1 mm, (D,L) = 500 µm, (C,E,H,K,J) = 250 µm, (P,I) = 100 µm.

By E21, the MOB undergoes remarkable maturation and achieves a laminated pattern comparable to that of the adult olfactory bulb. Six distinct layers can be clearly identified: the olfactory nerve layer, a single row of well-defined glomeruli surrounded by dense populations of periglomerular cells, the external plexiform layer, a well-formed mitral cell layer, the internal plexiform layer, and a deeply organized granule cell layer (Fig. 4G–I). This mature stratification strongly mirrors that of adult specimens of fossorial water voles. Medial to each MOB, vomeronasal nerve bundles are clearly visible, providing anatomical evidence of vomeronasal input reaching the AOB (Fig. 4I).

The AOB at E21 continues to display less advanced architecture. While a vomeronasal nerve layer is present and a cluster of principal cells with mitral-like features can be identified, glomeruli are not visible, and the surrounding plexiform zones remain poorly delineated (Fig. 4J–4K). Finally, a more caudal section illustrates the topographical relationship between the telencephalon, the olfactory peduncle, and the nasal cavity, offering broader spatial context for the understanding of the organization of the olfactory system at this stage (Fig. 4L).

### Immunohistochemical study

G proteins are essential components of the vomeronasal signal transduction cascade, and their detection may provide insight into the morphofunctional maturity of this system at early developmental stages. The expression patterns of the Gαi2 and Gαo subunits in fossorial water vole embryos was studied at stages E17 and E21 (Fig. 5). Immunohistochemical detection of Gαi2 (Fig. 5A–G) revealed moderate expression at E17 (Fig. 5A), with immunoreactivity present in both the sensory (SE) and respiratory (RE) epithelia of the VNO, being more pronounced in the SE. By E21 (Fig. 5B), this pattern intensified, with slightly stronger apical labeling in the SE, although without a clearly defined apicalbasal zonation. At this stage, Gαi2-positive axon bundles could be observed in the VNO (Fig. 5B) and in the vomeronasal nerve bundles (VN), coursing medially to the MOB. Remarkably, a subset of axons remained unlabeled with anti-Gαi2 (Fig. 5C). Transverse sections of the OB (Fig. 5D,E) showed weak Gαi2 labeling at E17, restricted to the superficial portion of the presumptive AOB. At E21, however, labeling was markedly stronger and confined to the AOB. Hematoxylin counterstained slides (Fig. 5F,G) confirmed immunoreactivity at E21 in both the VN layer (open arrowhead) and the glomerular layer (arrowhead).

**Figure 5.**
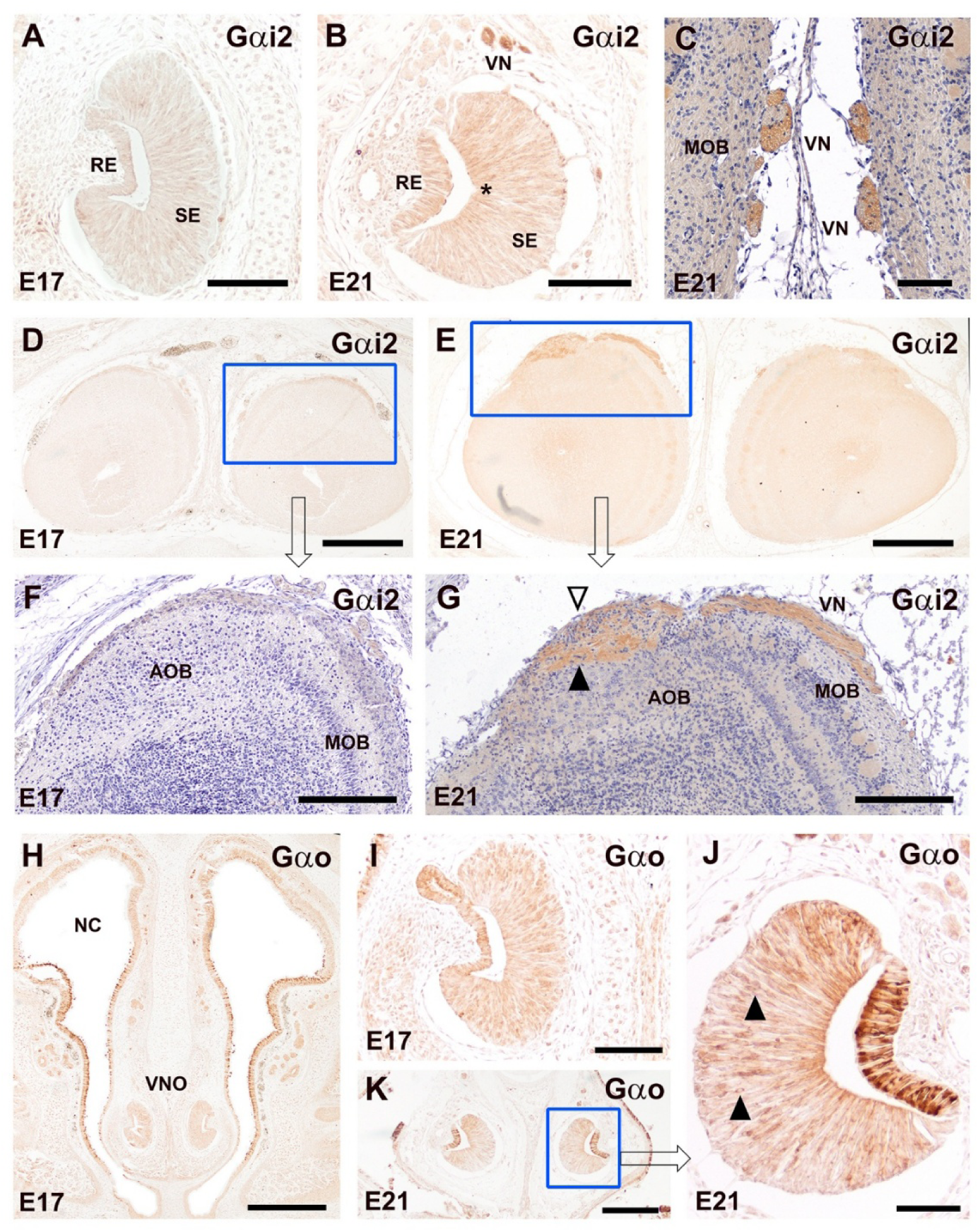
Immunohistochemical study of the G protein subunits αi2 and αo expression in the prenatal *Arvicola scherman*. **(A–G)** Immunolabeling with anti-Gαi2. (A,B) Transverse sections of the VNO at embryonic day E17 (A) and E21 (B), showing a progressive increase in Gαi2 expression in both the sensory (SE) and respiratory epithelium (RE). In the SE, immunoreactivity is more intense in the apical region, although no clear apical-basal zonation is observed. (C) In the vomeronasal nerves (VN) coursing along the medial surface of the MOB a subset of axons shows Gαi2-immunopositive labeling, though not all VN fibers are immunoreactive. (D,E) Transverse sections of the OB at E17 (D) and E21 (E). The AOB area is magnified atier counterstaining with hematoxylin (F, G, respectively). **(F)** At E17, only weak labeling is observed, restricted to the superficial layer of the AOB primordium. **(G)** By E21, intense Gαi2 immunoreactivity is present in the VN layer (arrowhead) and in the superficial glomerular layer (open arrowhead) of the AOB. The MOB remains negative for Gαi2. (H–K) Immunolabeling with anti-Gαo. **(H,I)** At E17, strong immunoreactivity is detected in the VNO and in the olfactory neuroepithelium. **(J,K)** At E21, this pattern persists, with a higher density of immunopositive somas in the basal layer of the SE (arrowheads), though no defined basal-apical zonation is apparent. Abbreviations: NC: nasal cavity. Scale bars: (D,E,H) = 500 µm, (F,G,K) = 250 µm, (A-C,I) = 100 µm, (K) = 50 µm.

Analysis of Gαo expression (Fig. 5H–K) revealed intense immunoreactivity in the VNO at E17 (Fig. 5H,I), with strong labeling throughout the sensory neuroepithelium. At E21 (Fig. 5J,K), Gαo immunoreactivity was still present, with stronger intensity in the basal portion of the SE, although no clear zonation pattern could be established.

The study of G protein subunits was extended to include the γ8 subunit (Gγ8). At an early stage as E17, strong Gγ8 immunoreactivity was detected throughout the VNO epithelium (Fig. 6A,B). Gγ8 was also expressed in the olfactory epithelium (OE) at this stage (Fig. 6A). At E21, transverse sections at the level of the MOB revealed that the VN bundles were uniformly Gγ8-positive (Fig. 6C). Immunoreactivity was evident throughout the superficial glomerular layer of the MOB, and no unlabeled axon subpopulations were identified, indicating a general expression of this subunit in the projecting vomeronasal axons. At this stage, weak immunoreactivity was observed in the OE, while a stronger signal was evident in the ONs (Fig. 6G), reflecting early establishment of olfactory connectivity. The analysis of the VNO at E21 (Fig.s 6D, E) showed that Gγ8 expression persisted, although the labeling intensity appeared slightly reduced compared to E17. Nonetheless, the signal remained distributed throughout the sensory epithelium (SE), suggesting the continued presence of Gγ8 in vomeronasal sensory neurons.

**Figure 6.**
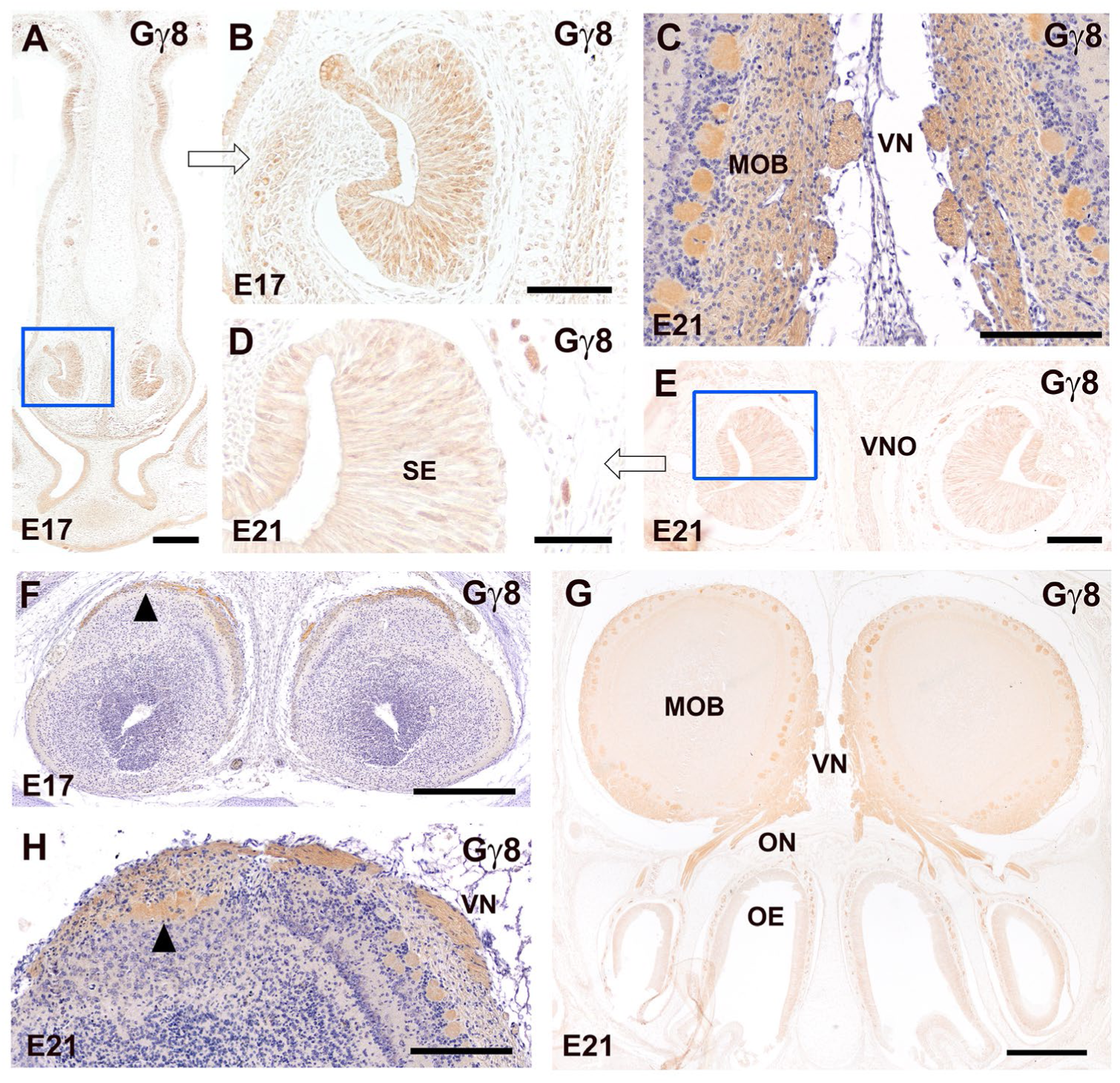
Immunohistochemical study of the expression of the G protein subunit Gγ8 during the prenatal development of the vomeronasal and olfactory systems in *Arvicola scherman*. **(A, B)** At E17, strong Gγ8 immunoreactivity is observed in the VNO epithelia as well as in the OE. **(C,G)** A transverse section at the level of the medial aspect of the main olfactory bulb (MOB) at E21 shows intense labeling in the vomeronasal nerve (VN) bundles and in the superficial glomerular layers of the MOB. No negative axon subsets can be distinguished, as all vomeronasal afferents appear Gγ8-immunopositive. Both the OE and the ON are strongly stained at this stage. **(D, E)** Examination of the VNO at E21 also reveals strong and widespread labeling across the sensory epithelium (SE), although slightly less intense than at E17. **(F)** At E17, the AOB shows superficial labeling restricted to its VN layer (arrowhead), with higher intensity than in the ON layer of the MOB. **(H)** At E21, the AOB shows intense Gγ8 immunoreactivity in both the VN and GlL layers (arrowhead), whereas in the MOB labeling is more concentrated in the ON and GlL layers. Scale bars: (G) = 500 µm, (A,C,F) = 250 µm, (B,E,H) = 100 µm, (D) = 50 µm.

In the OB at E17, Gγ8 expression in the AOB was limited to the superficial glomerular layer (Fig. 6F) and appeared more intense than in the MOB. At E21, the AOB exhibited intense Gγ8 immunoreactivity in both the VN and GlL layers (Fig. 6H), while the MOB showed a weaker labeling, mostly restricted to the nerve layer and with higher intensity in the glomerular region.

Immunohistochemical labeling with calcium binding proteins was employed to investigate the development of vomeronasal and olfactory systems. The study with anti-calbindin (CB) (Fig. 7) showed in counterstained coronal sections of the NC at E17 that CB expression was restricted to the VNOs (Fig. 7A). The immunolabeling extended across the SE and the central part of the NS epithelium (Fig.7B). Most positive neurons were confined to the basal third of the sensory layer, although some were organized into vertical columns projecting toward the apical surface. These structural arrangements were no longer evident at E21 (Fig.7C,D), when the immunoreactivity appeared more diffuse and was no longer detectable in the adjacent non-sensory epithelium. Regarding the AOB, at E17 only a subpopulation of vomeronasal sensory neurons expressed CB (Fig.7D). In the olfactory bulbs, E17 specimens showed intense immunoreactivity in the superficial layer of the AOB, particularly at the entry zone of the vomeronasal nerve (Fig.7E). Scattered CB-positive cells were also found within the MOB (Fig.7F).

**Figure 7.**
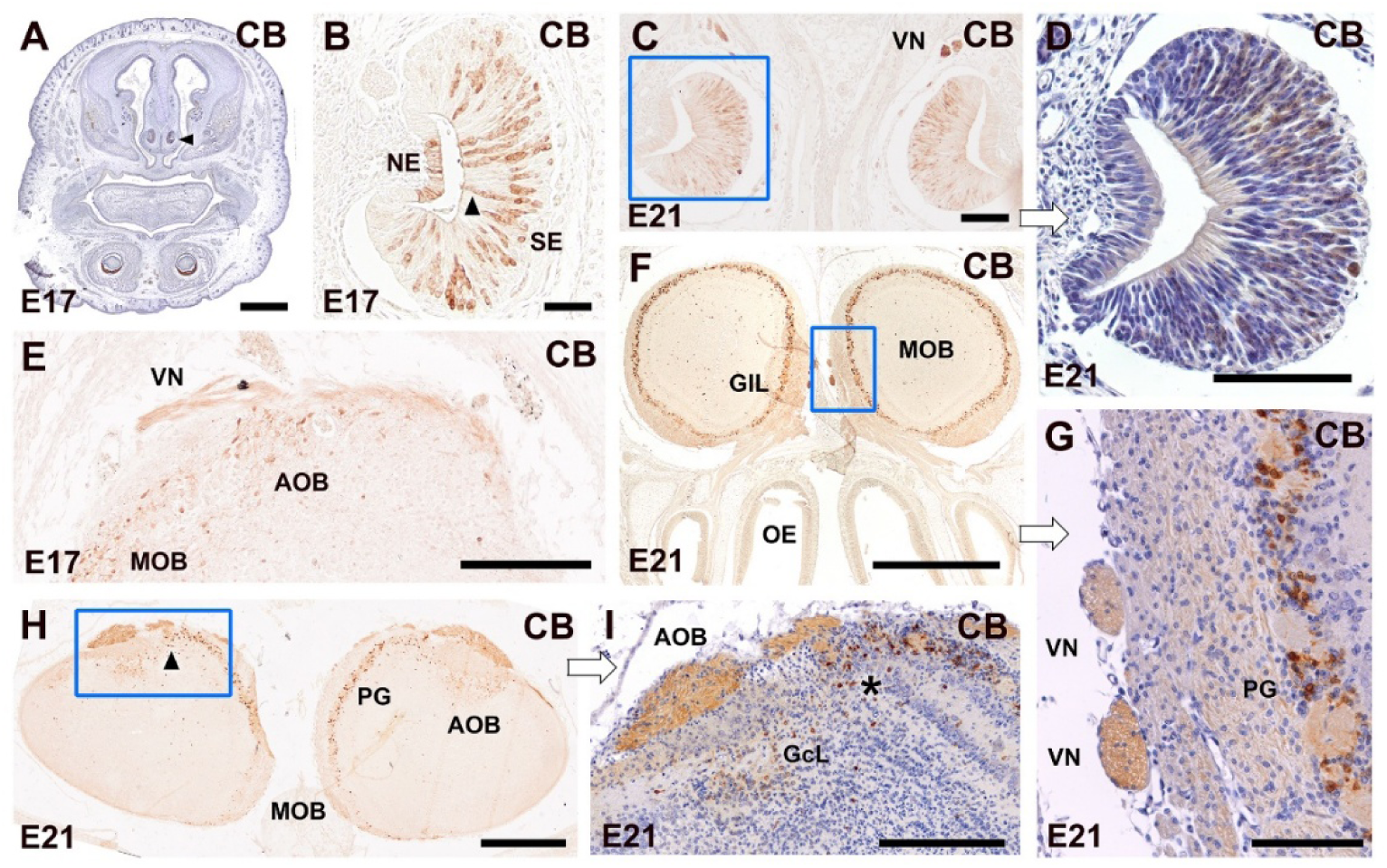
Immunohistochemical study with anti-calbindin (CB) of the vomeronasal and olfactory systems of the fossorial water vole. **(A)** Low magnification transverse section of the nasal cavity at E17 showing CB immunoreactivity restricted to the VNOs (arrowhead). **(B)** Higher magnification of a single VNO displaying CB immunoreactivity in both the sensory epithelium (SE) and the central region of the non-sensory epithelium (NE). Remarkably, most immunoreactive neurons are located in the basal portion of the sensory epithelium, with some columnar arrangements of immunopositive cells reaching the apical surface (arrowhead). **(C-D)** At E21, the labeling pattern in the VNO becomes more diffuse and the immunoreactivity in the NE is lost; columnar arrangements are no longer observed. **(E)** At E17, the labeling in the AOB is restricted to the superficial layer and the vomeronasal nerve (VN). Scattered immunoreactive cells are found in the MOB. **(F-G)** At E21, CN immunolabeling has become much stronger and widespread. In the MOB, the staining is concentrated in periglomerular cells **(PG; F,G)** and in the VN coursing medially to the MOB (G). **(H,I)** In the P21 AOB, strong CB expression is observed in the superficial nerve layer, although a distinct glomerular layer is not apparent. A high density of CB-positive cells is seen in the granule cell layer (GcL) of the AOB and in the PG cells of the MOB. The transition zone between the AOB and MOB also shows a higher density of CB immunoreactive cells (arrowhead in H, asterisk in I). Abbreviations: GlL: Glomerular layer. Scale bars: (A,F) = 1mm, (E) = 500 mm, (C,D,G-I) = 100 mm, (B) = 50 mm.

By E21, the anti-CB immunolabeling in the OB was neatly stronger. In the MOB, CB was expressed in periglomerular cells (Fig.7G) and in the medially located vomeronasal nerves (Fig.7F). In the AOB (Fig.7H), CB immunoreactivity was still observed in the superficial nerve layer, although no clearly differentiated glomerular structure could be identified. Interestingly, the granule cell layer of the AOB displayed a high density of CB-positive cells. The transitional zone between the AOB and MOB exhibited an increased density of CB-immunoreactive neurons, suggesting the presence of regionally enriched subpopulations.

Immunohistochemical analysis using anti-calretinin (CR) in prenatal fossorial water vole revealed a restricted and developmentally regulated pattern of expression within the nasal cavity and olfactory bulbs (Fig. 8). At both E17 and E21 stages, CR immunoreactivity was circumscribed to the sensory epithelia of the VNO and OE, with particularly strong labeling observed in the vomeronasal nerves and sensory neuroepithelial cells (Fig.8A,B). Remarkable, the OE and VNO remained the only immunopositive structures within the nasal cavity at both stages examined. At E17, CR expression was restricted to a subset of neuroepithelial cells, as observed in hematoxylin-counterstained sections (Fig.8C). By E21, this subpopulation expanded markedly, comprising morphologically diverse neuroepithelial cells, indicative of progressive neuronal differentiation (Fig.8D).

**Figure 8.**
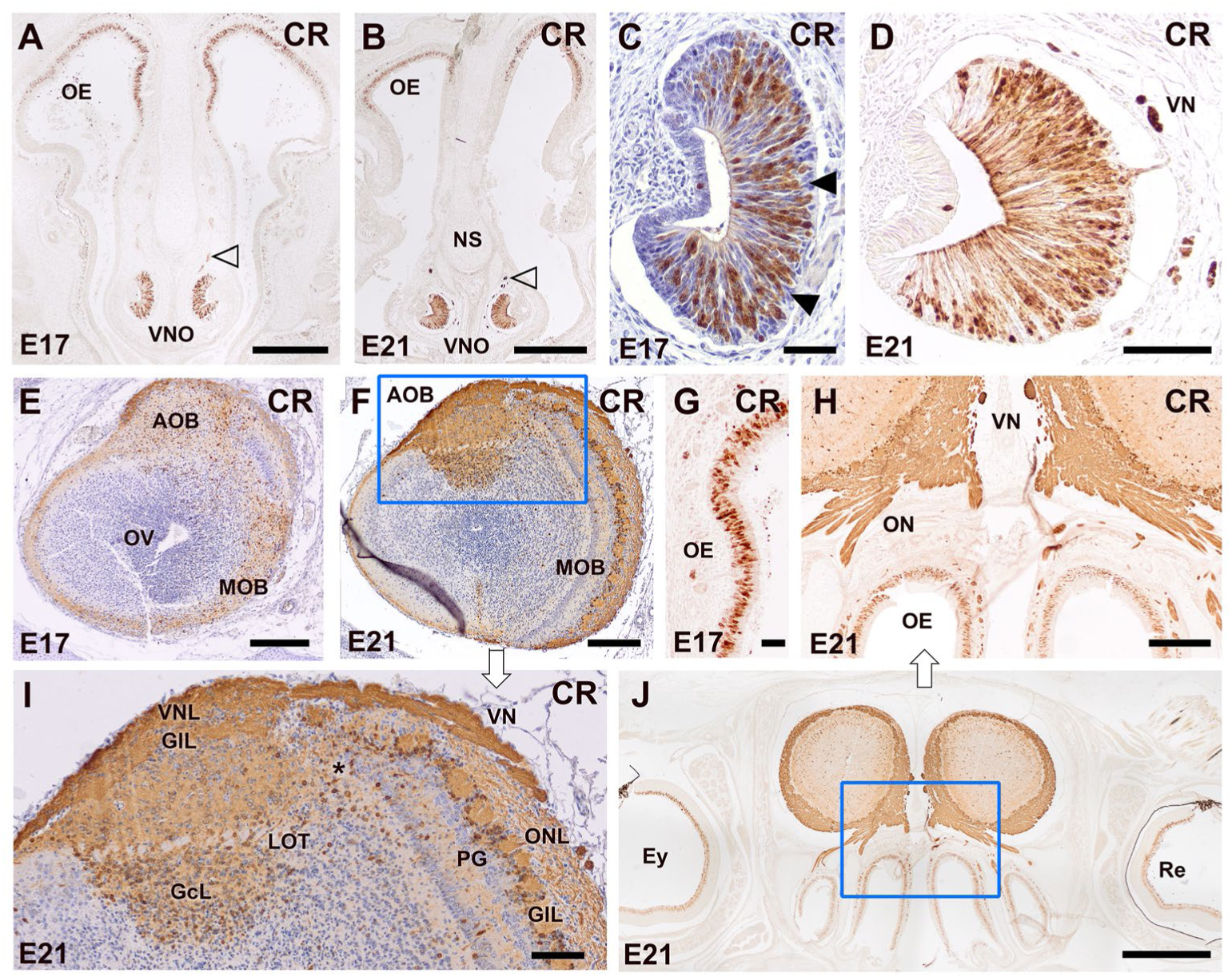
Immunohistochemical study with anti-calretinin (CR) of the vomeronasal and olfactory systems of the fossorial water vole. **(A)** Expression of CR in the vomeronasal organ (VNO) and olfactory epithelium (OE) shows that these are the only immunopositive structures within the nasal cavity at both embryonic day 17 (E17) and day 21 (E21). Immunolabeling intensity is strong in both the OE and the sensory epithelium of the VNO, including the vomeronasal nerves (arrowheads). **(B–C)** At higher magnification in CR-immunostained sections counterstained with hematoxylin, the E17 VNO neuroepithelium exhibits immunoreactivity restricted to a subpopulation of cells. **(D)** By E21, this subpopulation expands substantially, comprising neuroepithelial cells of diverse morphologies, likely at varying stages of differentiation **(E–F, I)** In the OB at E17, labeling is more intense in the area corresponding to the vomeronasal nerve layer (VNL) of the AOB, whereas the MOB and the remaining layers of the AOB show diffuse staining (E). By E21 (F), strong generalized CR immunoreactivity is observed throughout all layers of the AOB. At higher magnification (I), CR expression allows clear identification of the glomerular layer (GlL) and reveals a subpopulation of immunopositive cells in the mitral and plexiform layers, and especially in the granular cell layer (GcL). Fibers of the lateral olfactory tract (LOT) remain unstained. In the MOB, staining in the olfactory nerve layer (ONL) is less intense compared to the vomeronasal nerve layer (VNL). Diffuse immunoreactivity is present in the neuropil of the glomeruli (GlL) and the surrounding periglomerular cells (PG). Deeper layers of the MOB show less intense labeling in both neuropil and cell bodies compared to the equivalent layer of the AOB. Remarkably, the transition zone between MOB and AOB exhibits high cellular density (*). **(G–H,J)** In the OE of the ethmoturbinate region, intense CR immunoreactivity is observed in the neuroepithelial cells. At E17 (G), cells show variable morphologies, whereas by E21 (H,J), a more mature organization is evident. Labeling also extends intensely to the olfactory nerve bundles that will form the ONL of the MOB. Vomeronasal branches coursing medially to the MOB display very strong immunoreactivity (H). Scale bars: (I) = 1mm, (A,B,F) = 500 mm, (D,J) = 250 mm, (E,H) = 100 mm, (C,F) = 50 mm.

In the olfactory bulbs, CR labeling at E17 was concentrated in the vomeronasal nerve layer of the AOB, while the other layers of the AOB and the MOB exhibited diffuse immunoreactivity (Fig.8E). By E21, generalized and intense labeling was evident across all AOB layers, allowing precise identification of the glomerular (GlL), mitral, plexiform, and granular layers (GcL) (Fig.8F,I). The latest showed the strongest cellular density and immunoreactivity. In contrast, the LOT fibers remained CR-negative. In the MOB, the ONL displayed weaker staining in comparison to the VNL (Fig.8I). Diffuse immunoreactivity was present in the glomerular neuropil and periglomerular cells. Deeper layers exhibited a lower intensity of staining than the granular layer of the AOB. Remarkably, the transition zone between the MOB and AOB was marked by a high density of CR-positive cells.

In the ethmoturbinate region, the OE showed intense CR immunoreactivity (Figure 8G,H,J). At E17, neuroepithelial cells displayed variable morphology (Figure 8G), while at E21, cellular organization suggested advanced maturation (Figure 8H). Olfactory nerve bundles and vomeronasal nerve branches coursing medially to the MOB showed strong immunostaining.

PGP 9.5 is an excellent marker of differentiated neurons. In our preparations (Fig. 9), it allowed the visualization of immunopositive neuronal elements in both the VNO and the OE as early as E17 and in both sensory organs and the OBs at E21. At E17, strong PGP 9.5 immunoreactivity was already present in the VNO, the vomeronasal nerves, and the OE, as well as in associated nasal glands, such as the lateral and septal glands (Fig. 9A). A higher magnification of the OE at this stage confirmed intense expression in the neuronal layers of the epithelium (Fig. 9B). In transverse sections through the VNO, PGP 9.5 labeling was especially strong in the sensory neuroepithelium, where immunopositive cells frequently formed clusters or columns reaching the epithelial surface. This contrasted with the respiratory epithelium (RE), which showed a weaker and more diffuse pattern (Fig. 9C).

**Figure 9.**
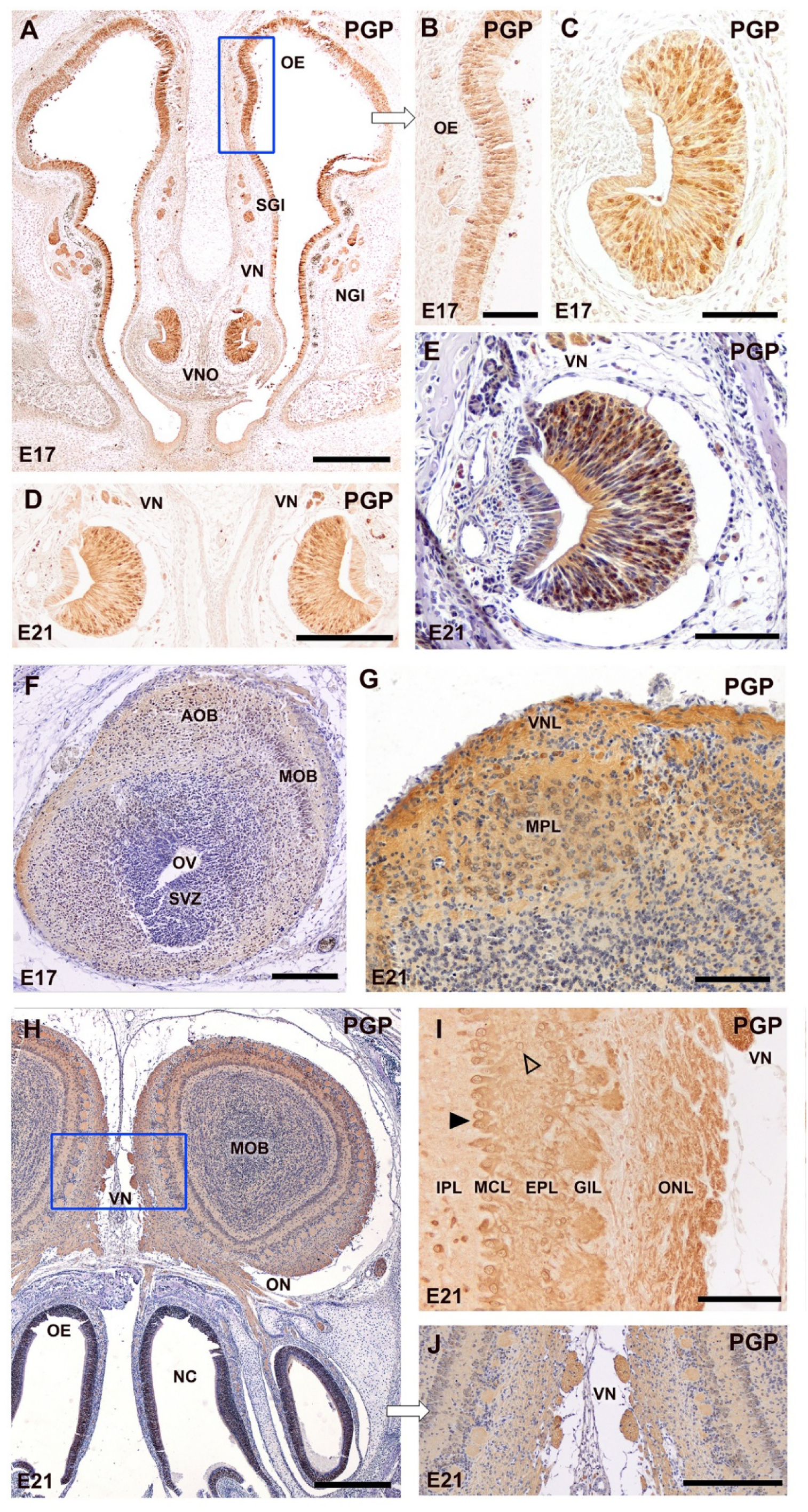
Immunohistochemical localization of PGP 9.5 in the vomeronasal system of the fetal fossorial water vole (A*. scherman*) at embryonic days 17 (E17) and 21 (E21). **(A)** Coronal section through the anterior nasal cavity at E17 showing strong PGP 9.5 immunoreactivity in the olfactory epithelium (OE), the vomeronasal nerves (VN), and the VNO. Strong labeling is also observed in the nasal septum glands (SGI) and the lateral nasal glands (NGI). (**B)** Higher magnification of the boxed area in (A), illustrating PGP 9.5 expression in the OE at E17. **(C)** Transverse section through the VNO at E17 showing strong PGP 9.5 immunolabeling in the sensory epithelium and surrounding nerve bundles. Labeling is more intense in neuroepithelial cells with a higher degree of differentiation, which otien cluster in groups or columns reaching the epithelial surface. In contrast, the respiratory epithelium displays weaker and less extensive immunoreactivity. (**D)** At E21, the labeling is more concentrated in the neuroepithelium and vomeronasal nerves (VN), and less intense in the glands and the RE. **(E)** In a hematoxylin-counterstained image at higher magnification, the labeling is observed to concentrate in the somata and processes of neuroepithelial cells, while sustentacular cells are not labeled. **(F)** Coronal section through the olfactory bulb at E17. The AOB is distinguishable from the MOB. The immunolabeling is very weak, limited to the superficial layers of the OB. **(G)** At E21, the AOB shows strong PGP 9.5 labeling in the vomeronasal nerve layer (VNL) and mitral plexiform layer (MPL). **(H)** Overview of the MOB at E21 showing intense labeling in all its layers and even stronger in the bundles of the VN (showed at higher magnification in (J). **(I)** At E21, PGP 9.5 immunolabeling allows visualization of a clear and well-defined lamination of the MOB, resembling the adult organization. A predominant presence of mitral cells (arrowhead), tutied cells (open arrowhead), and a clear distinction between the external plexiform layer (EPL) and the internal plexiform layer (IPL) can be observed. Abbreviations: NC: nasal cavity, ON: olfactory nerve, ONL: olfactory nerve layer, OV: olfactory ventricle, SVZ: subventricular zone. Scale bars: (A,H) = 500 µm, (D,F,G, J) = 250 µm. (B,C,E,I) = 100 µm.

At E21, PGP 9.5 immunoreactivity became stronger and more focused, concentrating within the vomeronasal sensory neuroepithelium and vomeronasal nerves, while diminishing in intensity in the surrounding glandular tissue and RE (Fig. 9D). Higher magnification revealed that labeling at this stage was confined to the somata and processes of neuroepithelial cells, with no signal detected in sustentacular cells (Fig. 9E).

In the OB, a developmental progression was also evident. At E17, both the AOB and MOB were distinguishable, but the overall immunolabeling was faint and limited to superficial layers (Fig. 9F). By E21, however, the AOB exhibited intense PGP 9.5 expression, particularly in the vomeronasal nerve layer and the mitral cell layer (Fig. 9G). Likewise, the MOB displayed strong labeling allowing visualization of a clear and well-defined lamination, resembling the adult configuration. Mitral and tufted cells and a distinct separation between the external and internal plexiform layers were evident (Fig. 9I). It is also remarkable the strong labeling at this stage of the vomeronasal nerve fiber bundles (Fig. 9H, 9J).

GAP-43 is a well-established marker of axonal growth and neuronal plasticity, particularly during early stages of neural development. Its expression is associated with axon elongation, synaptic remodeling, and differentiation processes. In the present study, GAP-43 immunoreactivity was used to evaluate the maturation of the vomeronasal and olfactory systems (Fig. 10). At E17, the VNO exhibited moderate GAP-43 immunoreactivity, mainly restricted to basal neuroepithelial cells, which appeared individually labeled, suggesting early neuronal differentiation within the sensory epithelium (Fig. 10A). By E21, labeling in the VNO became more diffuse and weaker, while also extending into the vomeronasal nerves (VN), indicative of ongoing axonal outgrowth (Fig. 10D).

**Figure 10.**
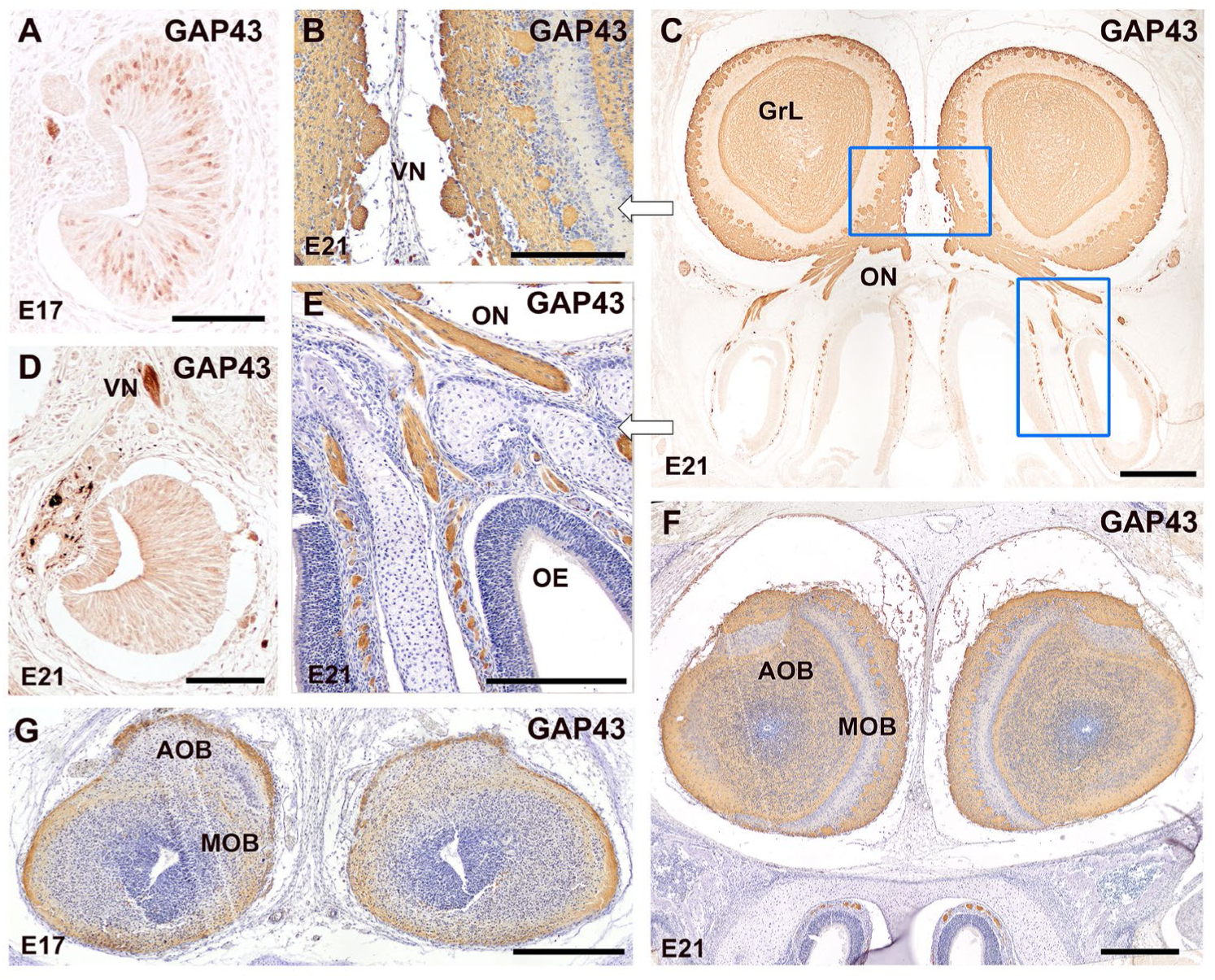
Immunohistochemical localization of GAP-43 in the vomeronasal and olfactory systems of the fetal fossorial water vole (*Arvicola scherman*) at E17 and E21. **(A)** Transverse section through the VNO at E17 showing GAP-43 immunoreactivity in the basal neuroepithelial cells, which are individually labeled. **(D)** At E21, GAP-43 expression in the VNO becomes more diffuse and less intense, extending to the vomeronasal nerves (VN). **(G)** In the main and accessory olfactory bulbs at E17, GAP-43 immunoreactivity is restricted to superficial layers, with labeling of the olfactory and vomeronasal nerve layers. **(C,F)** By E21, GAP-43 expression in the OB follows the mature pattern, with strong labeling of the glomerular layer (GrL) and outer nerve layers in both the MOB (C) and AOB (F), as well as moderate labeling in the granular layer. **(B,E)** Higher magnification of the boxed areas in (C) counterstained with hematoxylin confirms this pattern and shows intense GAP-43 immunoreactivity in the vomeronasal nerves coursing medially to the MOB (E). Strong immunolabeling is also observed in the ON emerging from the OE, indicating robust axonal growth. **(F)** The AOB exhibits a trilaminar pattern, with immunoreactivity in the superficial and deep layers, while the mitral-plexiform layer remains largely unlabeled. Similarly, the MOB shows a strong GAP-43 expression in the olfactory nerve and glomerular layers, and in the deeper granular zones, while the intermediate plexiform layers show reduced expression. Scale bars: (C,F,G) = 500 µm; (E) = 250 µm; (A,B,D) = 100 µm.

In the olfactory bulbs, both the MOB and AOB showed distinct labeling patterns at the two developmental stages. At E17, GAP-43 expression was confined to the superficial layers of both bulbs, with clear labeling of the olfactory and vomeronasal nerve layers, reflecting an immature organization (Fig. 10G). By E21, the pattern had shifted toward a more mature configuration. In the MOB, strong immunoreactivity was observed in the olfactory nerve and glomerular layers, with moderate labeling of the granular layer and lack of expression in the intermediate plexiform zone (Fig. 10C,F). A higher magnification view of the glomerular region (Fig. 10B) confirmed intense labeling of the VN projecting toward the AOB. Likewise, the ON emerging from the OE exhibited strong GAP-43 immunoreactivity as they left the nasal mucosa (Fig. 10E). The AOB displayed a clear trilaminar organization, with strong labeling in the superficial and deep layers, while the mitral-plexiform intermediate layer remained largely unlabeled (Fig. 10F). In contrast, the MOB exhibited a more continuous distribution, with strong immunoreactivity in the nerve and glomerular layers, diminished expression in the plexiform layers, and strong labeling reappearing in the deeper granular zones (Fig. 10F).

β-tubulin is a well-established marker of neuronal microtubules, and is widely used to assess the degree of neuronal differentiation, particularly in developing sensory epithelia (Fig. 11A-E). In the VNO, β-tubulin immunoreactivity was already prominent at E17, labeling a large population of neuroepithelial cells with well-defined neuronal morphology. These cells were distributed throughout the sensory epithelium, while sustentacular cells remained unlabeled (Fig. 11A). This pattern persisted at E21, with strong and widespread immunolabeling of the sensory VNO epithelium, suggesting that only a fraction of this epithelium is composed of differentiated neurons (Fig. 11B). A similar pattern was observed in the main olfactory epithelium, where β-tubulin strongly labeled olfactory receptor neurons and the associated axons in the lamina propria (Fig. 11C). Lower magnification images (Fig. 11D) revealed intense immunoreactivity extending from the OE through the ON bundles and reaching the β-tubulin positive MOB. Both the AOB and MOB showed widespread β-tubulin immunoreactivity as early as E17, labeling neuronal components across multiple layers (Fig. 11E).

**Figure 11.**
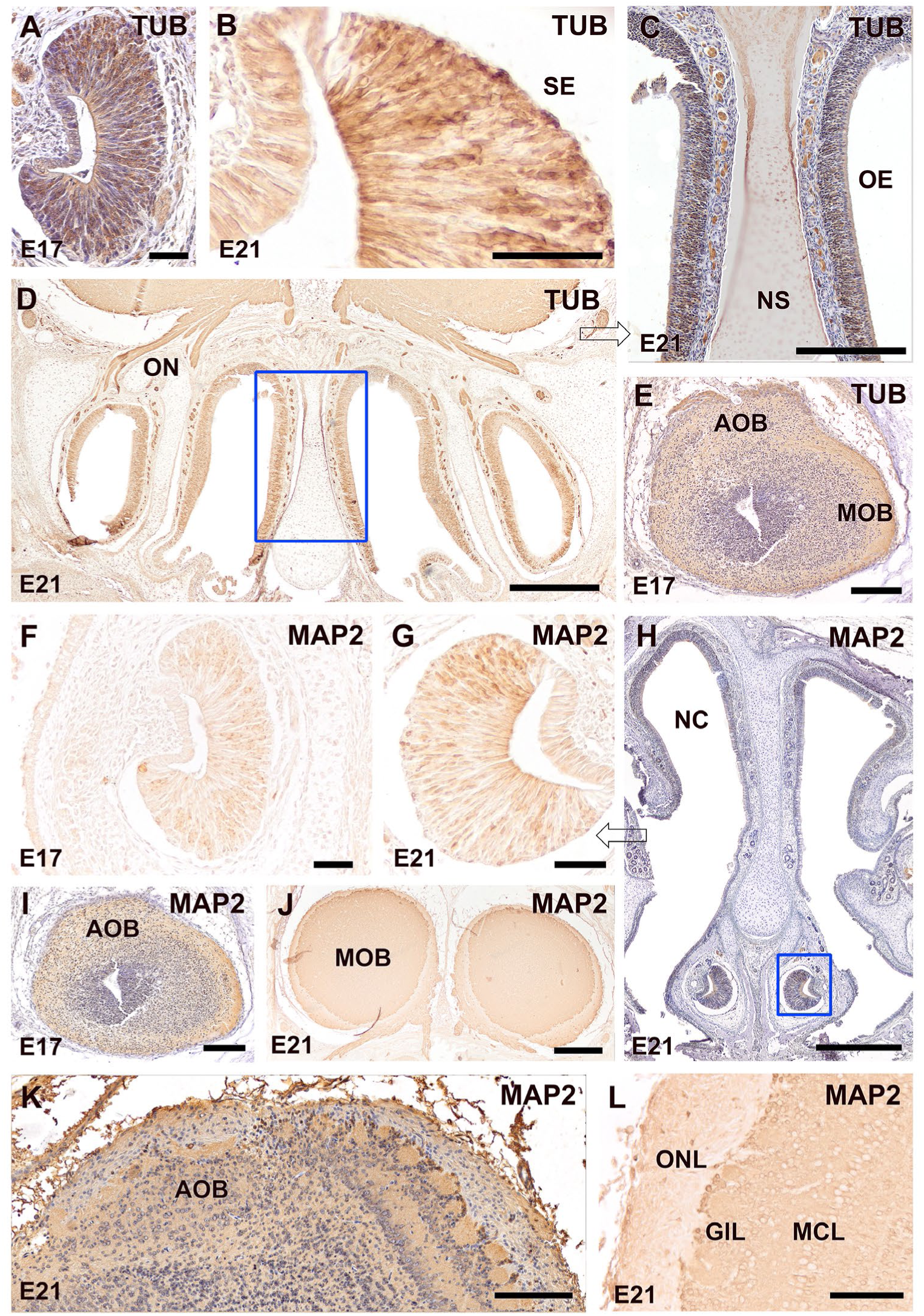
Immunohistochemical localization of β-tubulin (TUB) (A-E) and MAP2 (F-L) in the vomeronasal and olfactory systems of the fetal fossorial water vole (*Arvicola scherman*) at E17 and E21. **(A,B)** Transverse sections through the vomeronasal organ (VNO) showing intense TUB immunolabeling in a large population of neuroepithelial cells of the SE at both E17 (A) and E21 (B), with clear neuronal morphology and absence of staining in sustentacular cells. **(C)** The OE reveals at E21 strong TUB expression in the neuroepitelial cells and their axons in the lamina propria. **(D)** Low-magnification view of the nasal cavity at E21 shows TUB labeling throughout the sensory epithelia and nerve bundles which form ON which entry into the MOB. **(E)** At the level of the OB, TUB displays widespread expression in both the AOB and MOB already at E17, labeling neuronal components **(F–G)** MAP2 immunolabeling in the VNO shows faint signal at E17 (F), which increases markedly by E21 (G), highlighting neuroepithelial cells. **(H)** At E21, MAP2 expression in the olfactory epithelium remains weak overall, with minimal staining of the olfactory nerves and sensory epithelium. **(I–L)** In the OBs, MAP2 expression differs markedly from that of TUB. At E17 (I), a diffuse MAP2 signal is present in both the AOB and MOB. By E21 (J), labeling becomes restricted to deeper layers of the MOB and AOB, with no expression in the vomeronasal and olfactory nerve layers. (K) High magnification view of the AOB at E21 confirms this mature pattern, with diffuse labeling in deeper layers, from glomerular to granular, and scattered immunoreactive neuronal somata. (L) In the MOB, MAP2 labels the mitral cell layer (MCL) and granule cell layer (GCL) but spares the olfactory nerve layer (ONL). The presence of somatic labeling in superficial periglomerular cells is remarkable. Scale bars: (H) = 500 µm; (C-E) = 250 µm; (I,J) = 100 µm; (A,B,F,G) = 50 µm.

MAP2, in contrast, is a more selective marker of neuronal maturation, predominantly labeling dendritic processes (Fig. 11F-L). At E17, MAP2 expression in the VNO was weak and diffuse, indicating an early stage of neuroepithelial differentiation (Fig. 11F). By E21, MAP2 immunoreactivity increased substantially within the VNO sensory epithelium, though not comprising sustentacular cells, highlighting its specificity for neurons (Fig. 11G). However, MAP2 did not label vomeronasal nerves at either stage. In the olfactory epithelium, MAP2 immunoreactivity remained weak at E21, with minimal signal in the neuroepithelium and nerves (Fig. 11H). In the AOB, MAP2 showed diffuse labeling of deeper layers at E17 (Fig. 11I), and by E21, this pattern evolved into a more mature configuration. Expression was now restricted to the deeper regions of the bulb, with no labeling in the vomeronasal and olfactory nerve layers (Fig. 11J). Higher magnification views confirmed that MAP2 labeled both neuropil and scattered neuronal somata in the deep layers of the AOB (Fig. 11K).

In the MOB, MAP2 strongly labeled the mitral cell layer (MCL) and granule cell layer (GCL), while sparing the olfactory nerve layer (ONL) (Fig. 11L). Therefore, the immunolabeling observed in the glomerular layer is not attributable to the olfactory sensory axons themselves, which are MAP2-negative, but rather to dendritic arborizations of mitral and periglomerular cells projecting into this layer. Interestingly, somatic labeling was clearly observed in periglomerular neurons, which is unusual for a dendrite-associated marker like MAP2, and may reflect a late stage of perinatal maturation.

### Lectin histochemical study

UEA, a lectin with highest binding affinity for α-L-fucose residues, revealed distinct and dynamic patterns of glycosylation during the development of the vomeronasal and olfactory systems in *Arvicola scherman*. At E17, UEA binding in the VNO was restricted to discrete apical cells of the sensory epithelium, often grouped into columnar arrangements extending toward the luminal surface (Fig. 12A,B). This pattern shifted markedly by E21, with UEA reactivity strongly evident in the vomeronasal nerve bundles, as well as in the basal region of the sensory epithelium (Fig. 12C,D) Additionally, the respiratory epithelium showed moderate lectin binding at this stage (Fig. 12D).

**Figure 12.**
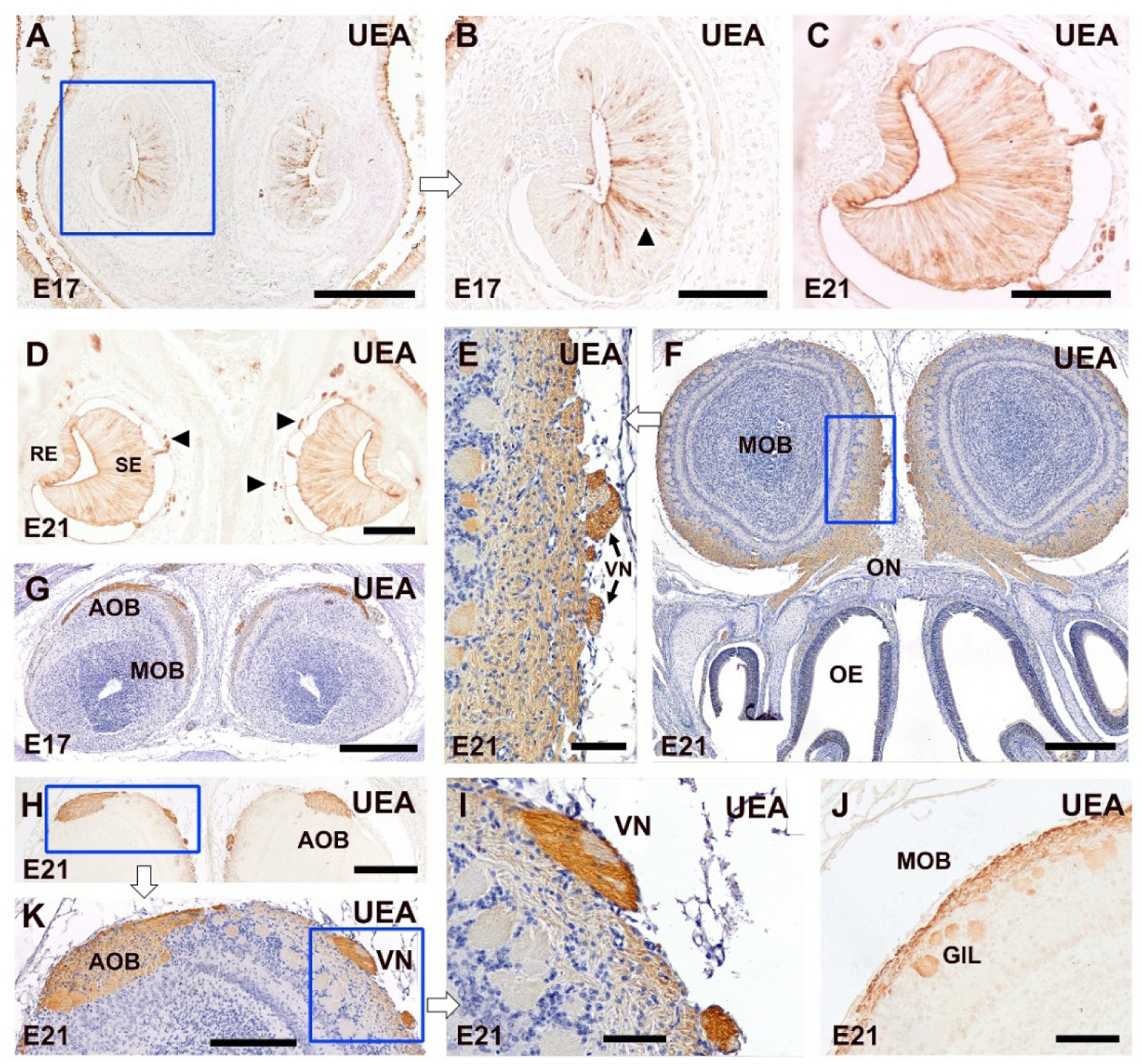
*Ulex europaeus* agglutinin (UEA) binding pattern in the vomeronasal and olfactory systems of the fetal fossorial water vole (*A. scherman*). **(A–D)** Transverse sections through the VNO at E17 (A,B) and E21 (C,D) showing changes in the UEA binding pattern during prenatal development. At E17 (A,B), UEA selectively binds to apically located cells in the vomeronasal SE, which otien form columns reaching the free surface (arrowhead). At E21 (C,D), UEA binding is markedly increased in the vomeronasal axons (arrowheads), and becomes concentrated in the basal region of the SE. Respiratory epithelium (RE) also shows moderate reactivity. **(E–J)** Transverse sections of the OB. At E17 (G), the AOB shows localized UEA positivity restricted to the superficial vomeronasal nerve layer. At E21, UEA binds to the olfactory nerve and glomerular layers of the MOB (E,F,J), whereas the olfactory epithelium (OE) remains largely negative (F). The branches of the vomeronasal nerve coursing medially to the MOB are strongly positive (E-K) Notably, labeling is stronger in peripheral axonal fibers (E,I), suggesting a functional segregation in the vomeronasal projections. Caudal transverse sections of the OB at E21 show strong UEA reactivity in the AOB nerve and glomerular layers (H,K). Scale bars: (F-H) = 500 μm; (A,K) = 250 μm; (B-E,I,J) = 100 μm.

In the OBs, at E17, UEA binding was restricted to the outermost nerve layer of the AOB (Fig. 12G). However, by E21, UEA labeling became more widespread, being consistently present throughout both the nerve and glomerular layers of the MOB (Fig. 12E,F,J), although the olfactory epithelium remained unreactive. In the AOB at E21, labeling was even stronger. The branches of the vomeronasal nerve projecting medially toward the bulb (Fig. 12E,I) were also strongly positive. Notably, only a subset of axons—typically those located peripherally within the nerve—were labeled, suggesting a potential functional segregation of vomeronasal projections within the AOB. Nevertheless, the AOB itself exhibited homogeneous and intense labeling across its outer layers, with no apparent zonation (Fig. 12H, K).

Both LEA and STA lectins, which exhibit high specificity for N-acetylglucosamine residues, revealed distinct patterns of glycosylation across the developing vomeronasal and olfactory systems (Fig. 13). At E17, LEA labeling was more prominent in the OE than in the VNO, where reactivity was limited to a few discrete apical cells (Fig. 13A, B). By E21, a marked increase in intensity and distribution was observed within the VNO, with labeling now predominantly located in the neuroepithelial layer (Fig. 13C). In the OB, both the main and accessory structures exhibited intense LEA binding within the nerve and glomerular layers at E17 and E21 (Fig. 13D-F,G). Olfactory nerve bundles were also strongly reactive. In contrast, STA staining was considerably weaker. Although it shares affinity for N-acetylglucosamine, STA showed only faint labeling in the VNO at both stages, again mostly in apical regions (Fig. 13G,H). In the OB, STA binding was also restricted to the superficial layers and showed low intensity compared to LEA (Fig. 13I,J).

**Figure 13.**
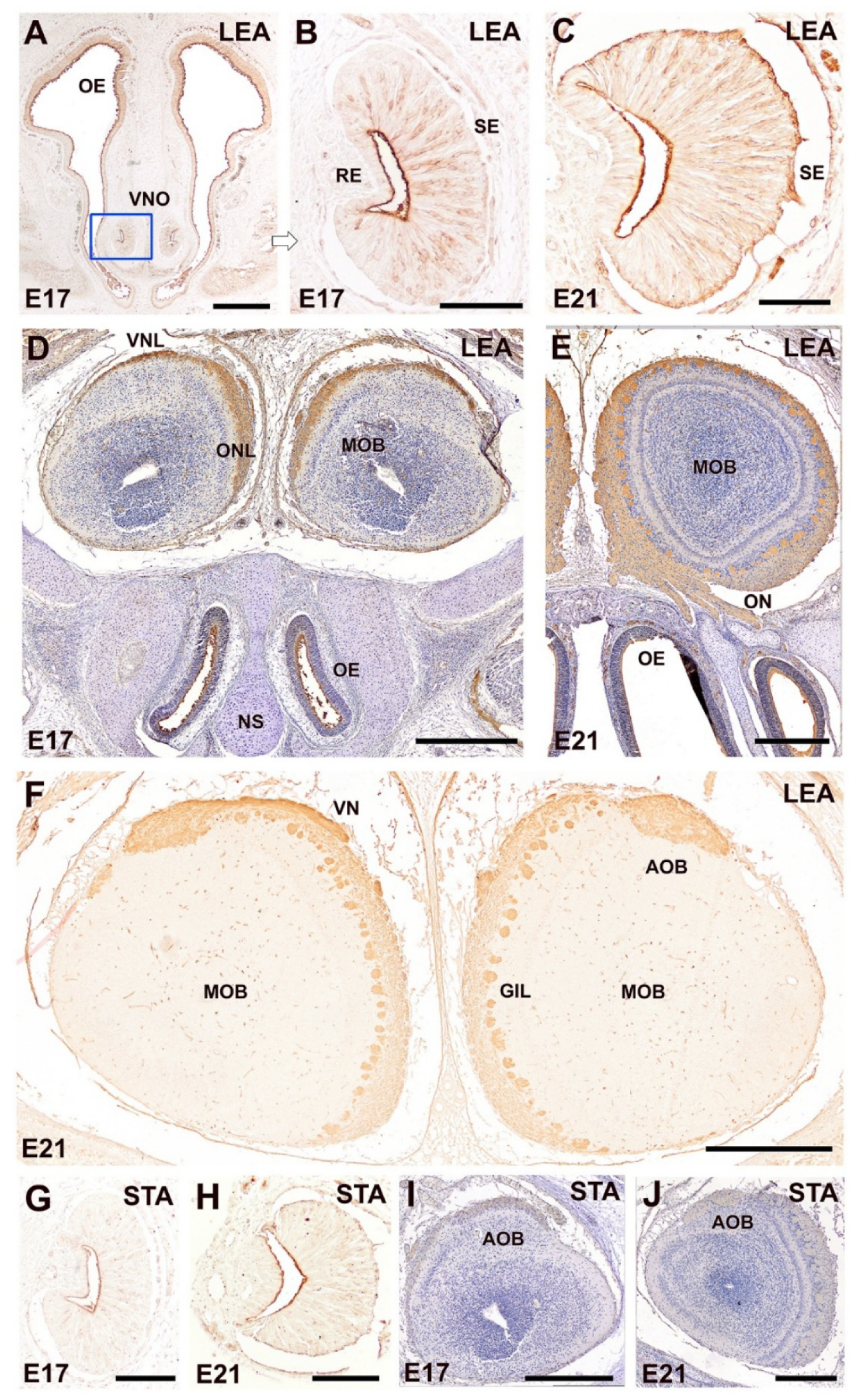
Histochemical detection of N-acetylglucosamine residues using LEA and STA lectins in the developing *A. scherman*. **(A–C)** Transverse sections of the VNO at E17 (A,B) and E21 (C) showing LEA binding patterns. At E17, the signal is mainly restricted to discrete apical cells within the vomeronasal SE, whereas the OE exhibits stronger reactivity. By E21 (C), LEA labeling becomes more widespread and intense in the VNO, particularly in the neuroepithelial cell layer. **(D–F)** Transverse sections of the OB stained with the LEA. At both E17 and E21, LEA shows strong and selective binding in the outermost layers—the nerve and glomerular layers—of both the MOB and AOB. Olfactory nerves are also intensely labeled (E). An image without hematoxylin counterstaining highlights the high specificity of LEA for the superficial layers. **(G–J)** STA lectin binding at E17 and E21. In the VNO (G, H), labeling is limited and restricted to apical regions, with only faint signal detected. Likewise, in the olfactory bulbs (I, J), STA staining is weak and confined to superficial layers. Scale bars: (A,D-F,I,J) = 500 µm; (B,C,G,H) = 100 µm.

The lectins SBA and DBA, both exhibiting high specificity for terminal α-N-acetylgalactosamine (GalNAc) residues, displayed distinct spatiotemporal binding patterns during the prenatal development of the vomeronasal and olfactory systems in *A. scherman*. SBA binding at E17 was weak and limited to the apical region of the vomeronasal sensory epithelium, with a small number of labeled neurons distinguishable (Fig. 14A,B). At this stage, the AOB exhibited weak SBA reactivity in the vomeronasal nerve layer and in incoming axons (Fig. 14C), while the MOB remained negative. At E21, a marked increase in SBA signal was observed in the AOB, which now showed intense labeling in both the vomeronasal nerve and glomerular layers, as well as in the vomeronasal projections approaching the bulb (Fig. 14E,F), indicating a selective affinity for components of the vomeronasal system.

**Figure 14.**
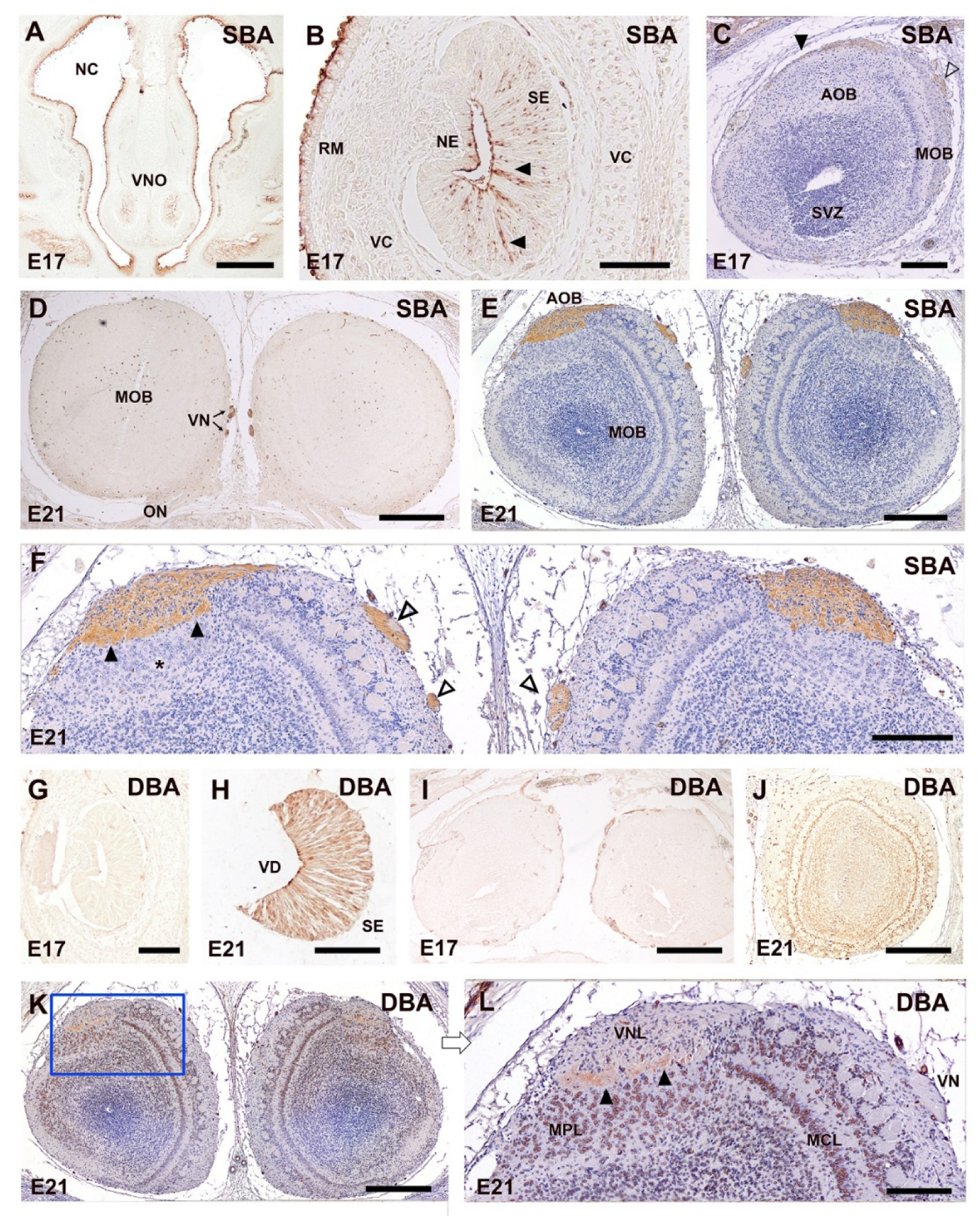
Histochemical detection of terminal α-N-acetylgalactosamine residues using SBA and DBA lectins in the fetal *A. scherman*. **(A–F)** SBA binding at E17 and E21. This lectin shows weak reactivity in the vomeronasal SE at E17, restricted to the apical region where isolated positive cells can be identified (arrowheads). In the AOB, weak labeling is observed in the superficial vomeronasal nerve layer (black arrowheads) and the incoming vomeronasal nerve (open arrowheads) (C). The MOB remains unstained. At E21, SBA binding is stronger and clearly specific for the vomeronasal system, labeling the vomeronasal nerves (D), and the vomeronasal nerve and glomerular layers (E,F) of the AOB. As observed with UEA, SBA binding in the vomeronasal nerve was restricted to a subpopulation of axons located peripherally within the fiber bundle, further suggesting a functional segregation of vomeronasal inputs within the AOB. **(G–L)** DBA binding at E17 and E21 shows negligible reactivity at E17 in both vomeronasal and olfactory structures (G-I). However, at E21, intense nuclear and cytoplasmic DBA binding is evident in periglomerular cells, mitral cells, and neurons within the mitral/plexiform layers of both the MOB and AOB (J-L), indicating late-stage expression of GalNAc-rich glycoproteins during olfactory system maturation. Scale bars: (A,D,E,I,J) = 500 µm; (C,F) = 250 µm; (B,G,H) = 100 µm.

In contrast, DBA binding showed a different temporal profile. At E17, DBA reactivity was virtually absent in both the vomeronasal organ and olfactory bulbs (Fig. 14G,I). However, by E21, intense DBA labeling emerged in both the MOB and AOB, with a clear nuclear and cytoplasmic pattern affecting periglomerular and mitral cells, as well as neurons in the mitral/plexiform layers (Fig. 14J-L). These findings suggest a late-stage appearance of GalNAc-rich glycoconjugates associated with neuronal maturation in both olfactory subsystems.

## DISCUSSION

Our prenatal study of the fossorial water vole (*Arvicola scherman*) shows a remarkably early and coordinated maturation of the nasal chemosensory systems. Across the E17–E21 interval—by E21, at term—the main olfactory bulb already exhibits adult-like lamination, the vomeronasal organ and olfactory epithelium display robust neuronal differentiation, and there is clear evidence of axonal growth and early circuit assembly before birth. In parallel, the accessory olfactory bulb lags behind the MOB in histologically evident glomerular organization, yet already exhibits selective molecular signatures in its nerve and superficial layers, indicating functional pathway specification despite incomplete morphogenesis. Together, these features indicate an accelerated functional readiness in a free-living, fossorial arvicoline and suggest ecological pressures favoring early chemosensory competence.

### Histological Analysis of Nissl-Stained Sections

By E21, the MOB exhibits a mature-like, six-layered architecture, with a single row of well-defined glomeruli, dense periglomerular populations, and sharply delineated mitral and granule layers—an organization closely resembling adults. In contrast, the AOB shows a vomeronasal nerve layer and a compact band of principal (mitral-like) cells but no identifiable glomeruli yet, indicating later maturation than the MOB. This level of definition of the fossorial water vole MOB at term is remarkable compared with laboratory mice: in BALB/c and C57 strains —as reported by Salazar et al. (2006) and Martin-Lopez et al. (2012), respectively—similar lamination and periglomerular profiles typically appear only postnatally (around P3–P5). Overall, the rodent pattern is preserved—MOB ahead of AOB—but in *A. scherman* the timetable is shifted forward, and molecular specification in the superficial compartments of the AOB precedes the appearance of morphologically distinct glomeruli.

At E17, relevant developmental features have already taken place in mice and rats: the first olfactory and vomeronasal axons have reached the anterior forebrain (Kim et al., 2023), the olfactory nerve is formed layer (Treloar et al., 2010), and the principal/mitral cell layer is evident (Treloar et al., 2003). Compared to the fossorial water vole, the interspecific differences at E17 are smaller: MOB axons have entered the presumptive glomerular zone and protoglomeruli are beginning to form; in the AOB, vomeronasal fascicles reach superficial layers, but glomerulogenesis remains very early.

In the periphery, the VNO of the fossorial water vole show at E17 and E21 conspicuous prenatal progression: clear stratification of the vomeronasal sensory epithelium, thick vomeronasal nerve bundles, and the onset of a bony capsule with vomeronasal glands evident by late gestation. Additionally, the VNO sensory epithelium shows prominent intraepithelial capillaries at E17, which become less abundant by E21. These morphological changes establish the anatomical substrate for prenatal chemical sampling and transport (Coppola and Millar, 1994), strengthening the case for early function. Comparable capillaries have been described in the rat VNO (Garrosa et al., 1986; Mendoza and Szabó, 1988; Szabó and Mendoza, 1988; Garrosa et al., 1998), but at similar prenatal ages they appear less developed than in *A. scherman*. In both fossorial water vole and rat, between E17 and E21, the shape of the VNO evolves from a latero-medially flattened profile to a more kidney-shape outline. The number and arrangement of layers in the sensory epithelium are broadly similar to rat, whereas the non-sensory epithelium in *A. scherman* tends to show a greater number of cell layers than reported for the rat by Garrosa et al. (1998). Within Cricetidae, the only other prenatal studies we identified concern the VNO of the golden hamster, *Mesocricetus auratus*, but they extend only to E15 (Taniguchi et al., 1982; Taniguchi and Taniguchi, 2008), preventing direct late-gestation comparisons with the fossorial water vole.

### Immunohistological Study

In the neuroreceptor cells of the adult mammals VNO, G-protein expression is a defining feature of the vomeronasal system (Halpern et al., 1995; Wekesa and Anholt, 1999). V1R neurons couple to the Gαi2 subunit and V2R neurons to Gαo (Halpern et al., 1995; Suárez et al., 2011), and are thus specifically involved in the signal transduction pathways of the two principal vomeronasal receptor families. Conversely to the αi2 and αo subunits, Gγ8 expression has been poorly studied in the vomeronasal system. This subunit is highly restricted to the sensory neuroepithelia of the OE and VNO, where must be playing a role in neuronal maturation/turnover in addition to signal transduction (Ryba and Tirindelli, 1995). Prenatally, available rodent data are scarce, but late-gestation expression in the olfactory and vomeronasal system has been documented in mice (Tirindelli and Ryba, 1996; Tirindelli, 2021). Because these markers delineate pathway identity, charting their appearance in the fetus is especially informative.

Prenatal immunohistochemical data in rodents on G-proteins expression are scarce. Beyond a specific report on Gγ8 from the Tirindelli group (Tirindelli and Ryba, 1996), systematic embryonic mapping of Gαi2/Gαo is largely missing. Our data indicate the onset of pathway specification and incipient connectivity of V1R/Gαi2 and V2R/Gαo subpopulations during late gestation, with Gγ8 highlighting broad afferent maturation prior to the emergence of morphologically distinct AOB glomeruli.

In particular, in the fetal VNO, both Gαo and Gαi2 are already detectable at E17, and their immunoreactive patterns become more prominent by E21. At both stages, Gαi2 labeling is more intense. Gγ8 displays a distinct profile, with strong expression in the VNO, the vomeronasal and olfactory nerves, and across the superficial layers of both the accessory and main olfactory bulbs at E17, followed by a slight reduction at E21. These results suggest that G-protein signaling is already involved in the early specification of the AOB, while Gγ8 may play a broader role in the maturation of vomeronasal circuits and their integration with olfactory processing areas.

Calcium-binding proteins such as calbindin, calretinin, and PGP 9.5 constitute a group of neuronal markers involved in intracellular calcium buffering, synaptic modulation, and neuronal differentiation (Baimbridge et al., 1992). These proteins have been widely used to characterize the emergence of interneuronal circuits, sensory projection pathways, and regional compartmentalization during prenatal development (Thompson et al., 1983; Jacobowitz and Winsky, 1991; Fujiwara et al., 1997). Their temporally and spatially regulated expression in the developing nervous system provides insight into the maturation status, connectivity, and functional specialization of distinct neuronal populations.

Calbindin is extensively expressed in adult neurons, particularly in GABAergic interneurons of the olfactory bulbs and hippocampus, where it contributes to calcium buffering and synaptic modulation (Celio, 1990). In prenatal stages, CB has been used as a marker of early postmitotic neurons and has been implicated in axonal targeting and regional specification in the olfactory bulb (Tufo et al., 2022). In our study, CB expression was already detectable at E17 in the vomeronasal sensory epithelium, with immunopositive cells primarily restricted to the basal third of the neuroepithelium and forming apically directed columns. This organization, suggestive of early differentiation and axonal guidance, disappeared by E21, when CB labeling became more diffuse. Interestingly, at E17 isolated immunopositivity was also detected in the non-sensory epithelium, but this signal vanished by E21, a finding consistent with the transient labeling observed with Gα0 in the same epithelial domain. In the olfactory bulbs, CB immunoreactivity intensified markedly between E17 and E21, especially in the superficial layers and granule cell layer of the AOB. At the cellular level, labeling was evident in granular cells of both the AOB and MOB; however, at the glomerular level, a strong and well-defined pattern was only present in the periglomerular cells of the MOB, while the AOB glomerular layer remained devoid of immunopositive periglomerular cells. This disparity further supports the delayed maturation of the AOB compared to the MOB, as the conventional histological organization suggested.

Calretinin (CR) is a calcium-binding protein related to calbindin but with more restricted expression patterns. In the adult olfactory bulbs, CR is enriched in periglomerular and superficial granule cells, and contributes to shaping inhibitory microcircuits (Kosaka and Kosaka, 2007; Torres et al., 2021). Its expression is associated with synaptogenesis and functional maturation (Schwaller, 2014). In the nasal cavity of prenatal *A. scherman*, CR expression was restricted to the sensory epithelia of the OE and VNO at both E17 and E21, with a marked increase in immunoreactive cell number and morphological complexity by E21. Within the OBs, CR labeling was initially mostly concentrated in the vomeronasal nerve layer of the AOB, but expanded by E21 to define distinct layers of the AOB, including the glomerular, mitral-plexiform, and granular layers, the latter showing the strongest intensity compared to E17. This pattern closely resembles that described in adult rodents (Porteros et al., 1995; Torres et al., 2020), suggesting the presence of a functionally mature circuitry despite the incomplete differentiation of glomeruli. This advanced organization not only highlights the early establishment of inhibitory microcircuits within the AOB. While in the E21 MOB, CR expression is primarily restricted to PG cells and a small subset of granular and mitral neurons, in the fossorial water vole, although no clear periglomerular labeling could be identified, numerous CR-positive principal neurons were found in the mitral and plexiform layers of the AOB, pointing to an advanced stage of mitral cell differentiation by this developmental period. Our observations suggest that the differences between the AOB and MOB extend beyond their distinct morphological maturation timelines: their developmental trajectories appear to follow divergent differentiation programs, which cannot be considered strictly analogous.

PGP 9.5 (Protein Gene Product 9.5), also known as UCHL1, is a ubiquitin hydrolase used as a general marker of postmitotic and differentiated neurons. In both adult and prenatal tissues, PGP 9.5 highlights the full extent of neuronal differentiation and axonal outgrowth (Thompson et al., 1983; Taniguchi et al., 1993a). It is particularly useful in sensory systems, including the olfactory epithelium and vomeronasal organ, where it labels sensory neurons and their projections (Kent and Rowe, 1992; Johnson et al., 1994). In prenatal studies, its pattern reflects the establishment of mature-like neural architecture. The study of PGP 9.5 expression in *A. scherman* is particularly relevant, since as early as E17 this marker shows widespread immunoreactivity. Unlike CB and CR, whose distribution was more restricted, PGP 9.5 was expressed not only in the olfactory and vomeronasal epithelia, but also extended to the respiratory mucosa and to the non-sensory epithelium of the VNO. The presence of immunopositive cells in this non-sensory epithelium is consistent with the transient labeling observed with the other calcium-binding proteins and with G-protein markers. However, by E21 the pattern had shifted from a diffuse organization to one resembling the adult, with clear labeling of vomeronasal neuroepithelial cells, while sustentacular cells were no longer immunopositive and labeling in the non-sensory epithelium markedly reduced. Interestingly, similar transient immunoreactivity of the respiratory epithelium was described in rats at E17 (Johnson et al., 1994), only vanishing in the neonatal period. These findings further suggest that the fossorial water vole follows an accelerated developmental timetable compared to the laboratory rat. The functional logic of PGP 9.5 expression in the vomeronasal lateral epithelium likely relates to its role in early differentiation of this non-sensory epithelium.

In the OBs, it is remarkable that at E17 only minimal labeling was observed in both AOB and MOB, whereas by E21 a generalized expression pattern encompassed virtually all cellular and laminar components of the AOB and MOB, consistent with an advanced stage of neuronal maturation. Particularly noteworthy is the ability of PGP 9.5 to clearly delineate periglomerular cells in the AOB, identifying these inhibitory elements with high specificity. These observations underscore the value of PGP 9.5 as an excellent marker for characterizing both structural and functional elements of the vomeronasal system.

GAP-43 is a membrane-associated phosphoprotein highly expressed in growing axons and growth cones, where it plays a critical role in axon elongation, pathfinding, and synaptic plasticity (Benowitz and Routtenberg, 1997; Holahan, 2017). The prenatal development of GAP-43 immunoreactivity has been mostly studied in early developmental stages, particularly to characterize the initial formation of olfactory axons arising from the olfactory placode (Pellier et al., 1994). However, there are fewer data on later fetal stages, and most developmental studies have focused in postnatal development. Verhaagen et al. (1989) described the postnatal progression of GAP-43 expression in the olfactory system of rats starting from P1. Their immunohistochemical results showed a high intensity of GAP-43 labeling in both the olfactory nerve and epithelium during the early postnatal period, especially at P1. However, the expression in the OE gradually declined: by postnatal week 3.5 the intensity had dropped substantially, and by the fifth week, GAP-43 staining was almost absent in the epithelium, reflecting the consolidation of olfactory projections and reduced axonal plasticity.

Interestingly, in *A. scherman*, the main olfactory system exhibits a markedly different temporal profile compared to laboratory rats. At E21, there is intense GAP-43 immunoreactivity along the olfactory nerves as they extend from the olfactory mucosa through the cribriform plate toward the OB. Within the bulb, the labeling comprises the olfactory afferent layers -olfactory and glomerular- and even extending into the deeper granule cell layer. In contrast, the olfactory epithelium itself displays only minimal immunolabeling—resembling the pattern described in the rat around postnatal week five, when GAP-43 expression has already declined substantially. This suggests that, in this fossorial species, both the morphofunctional maturation of the olfactory epithelium and the consolidation of its axonal projections could occur before birth. The postnatal decline in GAP-43 observed in the rat thus appears to take place intrauterinely in *A. scherman*, indicating an accelerated or shifted developmental timeline adapted to early sensory readiness.

At E17, GAP-43 is present in both the olfactory and vomeronasal nerves, as well as in their corresponding superficial layers within the bulbs; however, no expression is detected in deeper layers at this stage. The expression of GAP-43 also intensifies in the VNS from E17 to E2. The immunolabeling remains strong in the vomeronasal nerve and organ, and in the AOB it extends beyond the nerve layer to also encompass the glomerular and granule cell layers. Altogether, these observations demonstrate that the fossorial water vole exhibits a rapid maturation of axonal pathways and bulb lamination during the prenatal period.

β-tubulin and MAP2 are widely used neuronal markers with distinct roles in cytoskeletal organization and neurodifferentiation. β-tubulin, a major component of microtubules, is expressed early during neuronal differentiation (Roskams et al., 1998). MAP2, in contrast, is largely restricted to dendritic compartments and is often used to monitor dendritic arborization and neuronal maturation (Dehmelt and Halpain, 2005; Salazar et al., 2006). In A. scherman, β-tubulin immunoreactivity was already prominent at E17 in both the vomeronasal and olfactory sensory epithelia, labeling a broad population of differentiating neurons. This pattern persisted at E21, consistent with sustained neurogenic activity and early maturation. Immunoreactivity extended continuously from the VNO and the olfactory epithelium through the olfactory nerve bundles to both the main and accessory bulbs, where labeling encompassed multiple neuronal layers from the earliest stage examined.

MAP2 showed a markedly different pattern. At E17, MAP2 signal in the VNO was faint and diffuse, consistent with an early dendritic specification phase. By E21, immunoreactivity increased significantly in the vomeronasal epithelium, though the labeling remained confined to neuronal elements. In contrast, the olfactory epithelium showed only minimal MAP2 signal, even at E21, possibly indicating delayed dendritic development compared to the VNO. In both olfactory bulbs, MAP2 expression evolved from a diffuse labeling at E17 to a more stratified distribution by E21. Coinciding with the observations in mice by Salazar et al. (2006), in both the MOB and AOB, the labeling became restricted to deeper domains, sparing the nerve layers. Immunoreactivity in the GlL originated not from afferent axons—which are MAP2-negative—but from dendrites of mitral and periglomerular neurons. In both bulbs, the occasional somatic labeling observed in periglomerular cells may reflect late-stage neuronal maturation.

### Lectin histochemical study

Glycosylation constitutes a post-translational modification that regulates key aspects of neural development, including axonal guidance, synaptogenesis, and cell–cell recognition (Plendl and Sinowatz, 1998). In the olfactory and vomeronasal systems, the spatial and temporal distribution of specific glycoconjugates can be mapped using plant-derived lectins, which bind selectively to terminal carbohydrate residues. Lectin histochemistry is a powerful tool to investigate the functional compartmentalization of sensory pathways during development (Key, 1998; Treloar et al., 2010). Among the lectins tested for us in *A. scherman*, *Ulex europaeus* agglutinin (UEA), *Lycopersicon esculentum* agglutinin (LEA), and *Solanum tuberosum* agglutinin (STA) display high affinity for L-fucose and N-acetylglucosamine residues, respectively, while soybean agglutinin (SBA) and *Dolichos biflorus* agglutinin (DBA) are selective for terminal α-linked N-acetylgalactosamine (GalNAc). These sugar motifs are differentially expressed across developing sensory epithelia, nerve fibers, and olfactory bulb layers, providing molecular signatures of neuronal specification, maturation, and connectivity (Key and Akeson, 1993).

Although previous studies in rodents have reported lectin binding patterns in adult or late postnatal stages—often to delineate functional zones or glomerular subdomains—(Barber, 1989; Lundh et al., 1989; Pastor et al., 1992; Salazar et al., 2001; Salazar and Sánchez Quinteiro, 2003; Kondoh et al., 2017a; Keller et al., 2022) comprehensive developmental analyses remain scarce. The most extensive prenatal analysis in the olfactory systems of rodents is that of Franceschini et al. (1994) in the Wistar rat, who applied a panel of four lectins. Other contributions have focused on individual markers. Key and Akeson (1986) characterized SBA binding in both rat and mouse, while Plendl (1988) described DBA reactivity in the NMRI mouse. Later, Key & Akeson (1993) and Tisay et al. (2002) provided a detailed analysis of DBA patterns in the BALB/c mouse across prenatal and postnatal stages. More recently Lee et al. (2012) investigated the early postnatal development of the VNO in Sprague-Dawley rats with a panel of four lectins. These studies provided important references for the use of lectins in olfactory and vomeronasal development, but comprehensive and comparative analyses remain limited.

Among the lectins employed in this study, SBA proved the most selective marker for the VNS, without producing reactivity in the main olfactory pathway. Temporally, SBA expression appeared at term in E21 fetuses. At E17, SBA binding was weak but restricted to the apical portion of the vomeronasal neuroepithelium, consistent with early glycosylation events associated with vomeronasal. By E21, a remarkable increase in SBA labeling was observed in both the vomeronasal nerve bundles and the outer layers of the AOB, specifically the nerve and glomerular layers. This progression suggests a maturation-linked upregulation of GalNAc-rich glycoconjugates in vomeronasal projections and their target structures. Notably, the main olfactory bulb MOB remained completely negative at both developmental stages, reinforcing the utility of SBA as a specific marker of the prenatal vomeronasal pathway.

In the case of SBA, our results cannot be directly compared with other studies of prenatal development, as such data are lacking. However, information is available from studies in the adult olfactory and vomeronasal systems of mouse and rat. The study by Key & Giorgi (1986) confirmed that SBA is equally specific for the vomeronasal pathway in mice, with the peculiarity that in the rats it labels additionally a very limited subset of glomeruli. Additional evidence comes from studies in the adult mouse Keller et al. (2022), which reported SBA labeling not in the vomeronasal organ. At the level of the OB, only the MOB was examined, where SBA positivity was detected in the nerve and glomerular layers. It should be remarked, however, that in this study the tissue had undergone HIER pretreatment, a procedure known to alter lectin-binding profiles and likely responsible for the broader reactivity observed compared with our material. These observations point to a well-known feature of lectin histochemistry: the existence of considerable variability in labeling patterns, not only across species but also depending on methodological factors, including fixation and tissue processing (Takami et al., 1992; Lis and Sharon, 1998). Importantly, under the methodological conditions applied here—Bouin’s fixation, paraffin embedding, and without any pretreatment—SBA consistently emerged as a valuable and specific marker of the primary vomeronasal pathway in prenatal *A. scherman*.

In contrast, UEA, a lectin with high affinity for α-L-fucose residues, displayed strong labeling in both olfactory systems, although with distinct temporal and spatial patterns. At E17, UEA binding was restricted to discrete apical cells in the VNO and to the outermost nerve layer of the AOB, suggesting early compartmentalization of VNS projections. By E21, UEA reactivity became more widespread and intense, encompassing the sensory epithelium and the vomeronasal nerve fibers, and the outer layers of both the AOB and MOB. It is particularly striking that, with SBA, it becomes possible to distinguish within the population of vomeronasal axons coursing medially to the OB the presence of a negative subpopulation, i.e., a segregation between UEA-positive and UEA-negative fibers. This finding points to the potential establishment of a functional zonation in the vomeronasal system of the fossorial vole at this early developmental stage. The interpretation of this finding is further complicated by the fact that, in adult fossorial voles, the presence of such zonation in the AOB has not been observed consistently across individuals (Ruiz-Rubio et al., 2024a). In other mammal species, an anterior–posterior zonation has been reported, determined by the differential expression of G-proteins (Gαi2 and Gα0) as well as by UEA binding (Salazar and Sánchez Quinteiro, 1998; Salazar et al., 2006; Torres et al., 2022), although the exhaustive studies of Kondoh et al. in the mouse (2017a) demonstrated that UEA zonation in the adult AOB is not a constant feature. In our study we could not demonstrate such segregation within the prenatal fossorial water vole AOB itself. Nevertheless, it seems reasonable to interpret that most of the SBA-negative fibers correspond to Gα0-positive axons. On the other hand, at E21 it was not possible to determine a clear apical–basal zonation in the VNO of the fossorial water vole—nor in the adult (Ruiz-Rubio et al., 2024b)—unlike what has been reported in other rodents. This makes it more difficult to confirm our interpretation that UEA-negative fibers correspond to Gα0-positive axons. Finally, the postnatal development of the VNO in Sprague–Dawley rats, examined with the lectin UEA, showed no positivity at P1, with reactivity first appearing at P7 (Lee et al., 2012). This finding once again suggests that the developmental onset of glycoconjugate expression in *A. Scherman* occurs earlier than in laboratory rodents, reflecting a comparatively accelerated maturation process.

The group of LEA and STA, both N-acetylglucosamine-specific lectins, showed overlapping but quantitatively divergent reactivity. LEA was a reliable marker for both vomeronasal and olfactory structures, with moderate labeling of apical VNO cells at E17 and intense signal in the neuroepithelium as well as the entire vomeronasal nerve branches by E21. In the OBs, LEA labeled the nerve and glomerular layers of both MOB and AOB consistently from E17, confirming its utility as a pan-chemosensory marker. STA, however, showed a much weaker and more restricted pattern, with faint apical labeling in the VNO and only low-intensity staining in the OBs. This discrepancy, despite shared sugar specificity, suggests subtle differences in binding thresholds.

The binding pattern observed with LEA can be regarded as virtually pathognomonic of the olfactory and vomeronasal system across all mammalian species in which it has been investigated. Although developmental data are still lacking for the earliest prenatal stages, the distribution we observed is fully consistent with that reported in every species studied so far, where LEA reactivity invariably comprises both olfactory subsystems (Taniguchi et al., 1993b; Halpern et al., 1998; Salazar et al., 1998; Ibrahim et al., 2013; Tomiyasu et al., 2018; Jang et al., 2021). This high degree of conservation is unique among the panel of lectins usually employed in the studies of the nervous system, most of which typically display broad interspecific variability in both intensity and localization.

Finally, DBA revealed a unique temporal profile, characterized by late-onset labeling and prominent cellular reactivity. At E17, DBA signal was nearly absent in all regions. However, by E21, intense cytoplasmic and nuclear labeling appeared in periglomerular and mitral cells of both MOB and AOB, as well as in deeper neuronal layers. This pattern suggests that GalNAc-rich glycoconjugates targeted by DBA are linked to neuronal differentiation events occurring after the establishment of primary axonal projections. In this context, DBA may act as a marker of postmitotic maturation in projection neurons, mitral, tufted and principal cells.

Interestingly, our findings parallel—but also extend—those of Brian Key’s group, who carried out an exhaustive lectin study in the olfactory and vomeronasal pathways of postnatal BALB/c mice (Tisay et al., 2002). At E16.5, these authors described a weak and superficial DBA signal in the OB, closely resembling the limited reactivity we observed at E17 in *A. scherman*. By P0.5, they reported broader labeling involving postmitotic elements, particularly mitral cells, although the intensity remained modest compared with our observations in the fossorial vole at E21. Moreover, a striking qualitative divergence emerges: whereas in BALB/c mice DBA postmitotic mitral neurons were not found in the AOB, in *A. scherman* we found a clear and intense reactivity in the mitral/plexiform layer of the AOB at E21. This difference may indicate species-specific regulation of GalNAc-rich glycoconjugates during late prenatal development, potentially linked to ecological and evolutionary adaptations. To these studies can be added those performed in BALB/c (Salazar and Sánchez Quinteiro, 2003) and NMRI mice (Plendl and Schmahl, 1988), which detected lectin reactivity only in the AOB at the first postnatal day. Interestingly, the study of Lee et al. (2012) on the VNO of Sprague–Dawley rats reported DBA positivity only at eight weeks of postnatal life.

Together, these results underscore the value of combining multiple lectins to track the temporal emergence and compartmental specialization of glycosylation signatures during chemosensory development. SBA and UEA provide complementary information on the VNS, while LEA offers broader coverage of both main accessory olfactory pathways. DBA, although less specific, may provide valuable insights into the development and characterization of the secondary neurons of both olfactory pathways.

In conclusion, our prenatal study of the fossorial water vole demonstrates that the nasal chemosensory systems undergo an unusually rapid and coordinated maturation before birth. By term (E21), the MOB already exhibits an adult-like lamination, the vomeronasal and olfactory epithelia show robust neuronal differentiation, and multiple molecular markers—including G-proteins, calcium-binding proteins, and lectins—reveal the early specification of functional pathways. Although the AOB lags behind the MOB in its morphological glomerular development, the presence of distinct molecular signatures in its superficial layers underscores that functional organization precedes full structural maturation. Lectin histochemistry in particular, shows both conserved and species-specific glycosylation patterns: SBA confirms its role as a highly selective marker of the vomeronasal pathway; UEA reveals early compartmentalization of vomeronasal projections; LEA maintains a virtually pathognomonic profile across mammalian taxa; and DBA, although less specific, provides unique information on the maturation of bulbar projection neurons. Altogether, these findings suggest that *A. scherman* achieves prenatal chemosensory readiness through an accelerated developmental program compared with laboratory rodents, likely reflecting ecological pressures that favor early functional competence in fossorial environments.

## AUTHORS CONTRIBUTION

Sara RUIZ-RUBIO: Conceptualization; Investigation; Funding acquisition; Writing - original draft. Irene ORTIZ-LEAL: Conceptualization; Investigation; Writing - review & editing; Methodology; Supervision. Mateo V. TORRES: Conceptualization; Investigation; Methodology; Aitor SOMOANO: Investigation; Methodology; Supervision; Writing - review & editing. Taekyun SHIN: Investigation; Methodology; Supervision; Writing - review & editing. Pablo SANCHEZ-QUINTEIRO: Conceptualization; Investigation; Funding acquisition; Writing - original draft; Methodology; Writing - review & editing; Supervision.

## COMPLIANCE OF ETHICAL STANDARDS

## Ethical approval

Fossorial water voles are considered a key pest species in grasslands, and their demographic densities need to be controlled in accordance with article 15 of the Law 43/2002 of plant health (BOE 2008) and therefore the practices undertaken in this study are considered as a recognised zootechnical purpose (Real Decreto 53/2013). Accordingly, ethics approval was not required for this study. The recommendations of the Directive of the European Parliament and the Council on the Protection of Animals Used for Scientific Purposes (Directive 2010/63/UE 2010) were considered in all procedures.

## Informed consent

No human subject was used in this study.

## ACKNOWLEDGEMENTS

The authors wish to thank the “Dirección Xeral de Gandaría, Agricultura e Industrias Agroalimentarias of the Consellería do Medio Rural of the XUNTA de GALICIA” for the financial, logistical support, and the trust placed in this project.

